# Adolescent alcohol binge-drinking induces delayed appearance of behavioral defects in mice

**DOI:** 10.1101/2020.08.11.245878

**Authors:** Laura Van Hees, Vincent Didone, Manon Charlet-Briart, Théo Van Ingelgom, Alysson Alexandre, Etienne Quertemont, Laurent Nguyen, Sophie Laguesse

## Abstract

Adolescence is a developmental period characterized by significant changes in brain architecture and behavior. The immaturity of the adolescent brain is associated with heightened vulnerability to exogenous agents, including alcohol. Alcohol is the most consumed drug among teenagers, and binge-drinking during adolescence is a major public health concern. Studies have suggested that adolescent alcohol exposure (AAE) may interfere with the maturation of frontal brain regions and lead to long-lasting behavioral consequences. In this study, we used a mouse model of AAE in which adolescent mice reach high blood alcohol concentration after voluntary binge-drinking. In order to assess short- and long-term consequences of AAE, a battery of behavioral tests was performed during late adolescence and during adulthood. We showed that AAE had no short-term effect on young mice behavior but rather increased anxiety- and depressive-like behaviors, as well as alcohol consumption during adulthood. Moreover, alcohol binge-drinking during adolescence dramatically decreased recognition memory performances and behavioral flexibility in both adult males and females. Furthermore, we showed that voluntary consumption of alcohol during adolescence did not trigger any major activation of the innate immune system in the prefrontal cortex (PFC). Together, our data suggest that voluntary alcohol binge-drinking in adolescent mice induces a delayed appearance of behavioral impairments in adulthood.

## Introduction

Adolescence is a crucial developmental phase highly conserved across mammalian species, and typically defined as a transitional period between childhood and adulthood. This transition period involves significant changes in brain architecture, including cortical gray matter volume decline via synaptic pruning and increased white matter volume due to continued myelination of axons ^1^. Adolescence is also characterized by complex developmental changes in neural processing systems and unique behavioral characteristics including increased impulsivity, novelty-seeking and desire of risk-taking ^2,3^. Brain maturation typically begins in posterior brain regions and continues towards more anterior higher-order regions until ~25 years old ^1^. The prefrontal cortex (PFC) which is implicated in executive functions ^4^, is one of the last brain region to become fully mature, and immaturity of this brain region in adolescents is associated with lack of inhibitory control over behaviors ^2,3^. Moreover, adolescence is typically the age for initial exposure to a number of potentially toxic exogenous agents ^5,6^. Alcohol is the most consumed addictive substance among teenagers, with 27% of adolescents worldwide reporting alcohol consumption during the past month ^7^. Binge-drinking of alcohol, which corresponds to ingestion of at least five drinks in males (four in females) within a 2-hour period, has become a common pattern of alcohol consumption among teenagers. Binge-drinking leads to high blood alcohol concentration (above 0.08g/dl) ^8,9^, which can be harmful to adolescent brain, as it may interfere with ongoing maturation of its frontal circuits. Clinical studies reported that adolescent alcohol exposure (AAE) is associated with brain structure changes, comorbid psychopathology and detrimental neurocognitive consequences ^5,10,11^. Indeed, binge-drinking in adolescent has been associated with thinner cortical and subcortical structures, including the prefrontal cortex, and reduced white matter development ^12^. AAE is also believed to have deleterious effects on verbal learning and memory, attentional and executive functions ^11,13^, as well as to increase the risk of developing psychiatric and behavioral disorders later in life, including alcohol addiction ^5,10,14,15^. Altogether, it has become clear that, because of high brain plasticity, adolescence is a sensitive period for the development of alcohol-related behavioral impairments. Over the past years, rodent models have been used to study alcohol’s impact on the adolescent brain and have provided findings consistent with human research ^11,15,16^. Indeed, several studies demonstrated short- and long-term defects in executive functions and behaviors induced by AAE in rodents ^11,15^. Interestingly, most studies used a “binge-drinking-like” administration of alcohol, involving repeated i.p. injections or gavage, and showed that AAE induced activation of the innate immune system in frontal cortical regions ^17–20^ as well as short- and long-term behavioral defects. Here, we used a model of voluntary alcohol consumption during early/mid adolescence in mice and we examined the short- and long-term consequences of AAE on a battery of behavioral tests requiring the PFC. We reported that while not impacting adolescent behaviors, AAE strongly impairs behaviors during adulthood, similarly in males and females, without inducing major neuro-inflammation in the PFC.

## Material and Methods

Detailed information can be found in supplementary material.

### Animals

Males and females C57BL/6J (Janvier Labs, Saint Berthevin, France) were treated according to the guidelines of the Belgian Ministry of Agriculture in agreement with the European Community Laboratory Animal Care and Use Regulations (86/609/EEC) for care and use of laboratory animals under the supervision of authorized investigators (ethical file 18-2004). Experimental animals were bred in-house and maintained with *ad libitum* access to food and constant temperature (19° to 22°C) and humidity (40 to 50%) under a reversed light/dark cycle (lights on at 22:00, off at 10:00).

### Adolescent Alcohol Exposure

Adolescent males and females underwent a modified version of the Drinking in the Dark paradigm ^21^ from P29 to P33, and from P36 to P40 (Fig. 1E). The age for AAE was chosen during early/mid adolescence ^22^. All mice were group-housed in order to avoid social isolation stress. At 9:30, mice were weighted and transferred to single cages with water and food *ad libitum*. At 12:00, water was replaced by ethanol (20% in tap water). At 16:00, alcohol was removed and mice were group-housed until the next day. Control mice received only water and alternated between single- and group-housing accordingly. Different cohorts of mice underwent the behavioral tests either 72 hours after the last drinking session (P43, middle adolescence), or after 40 days of abstinence (P80, adulthood). Ninety-one percent of the animals drank more than 4g/kg/4h and were included in the study. Animals which drank less than 4g/kg/4h for more than 2 sessions were excluded. The threshold of 4g/kg/4h was chosen because it represents the amount of alcohol ingested leading to minimal binge-drinking BAC values, as previously described ^23^. Individual drinking data can be found in Tables S1 and S2.

**Fig 1:**
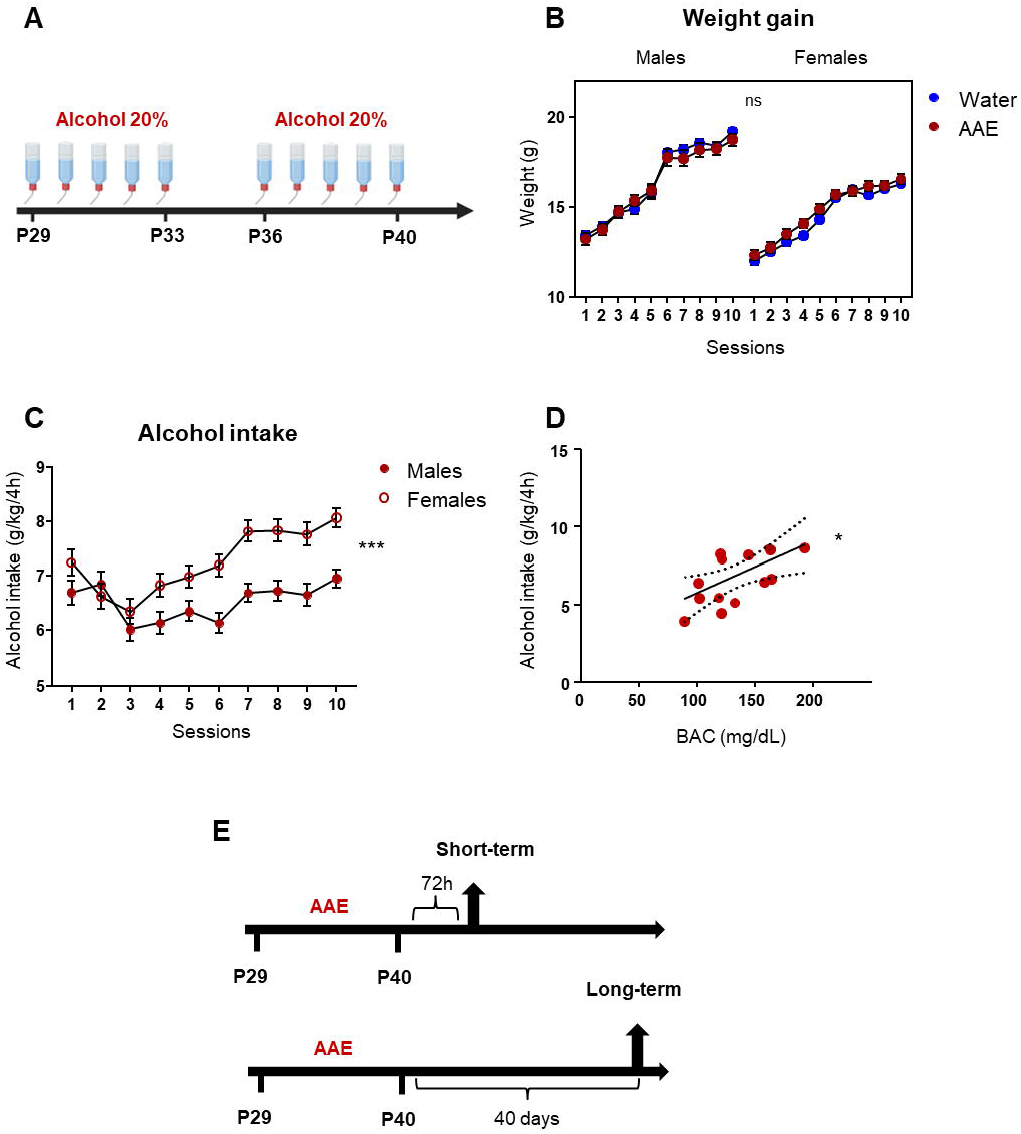
Adolescent mice voluntarily consume high amounts of alcohol and reach binge-drinking-related blood alcohol concentrations. **(A)** Model of alcohol exposure. Adolescent males and females have access to alcohol 20% for 4 hours per day, for ten sessions (from P29 to P33, and from P36 to P40). **(B)** Weight gain was measured over the course of the treatment and compared with water control littermates. Linear mixed-effects model revealed a main effect of session (χ^2^_(9)_=1.18×10^4^, p<0.001 η^2^=0.65) and sex (χ^2^_(1)_=51.41, p<0.001, η^2^=0.36), but no effect of treatment (χ^2^_(1)_=0.11, p=0.74); n=23 per group. **(C)** Voluntary alcohol intake. Linear mixed-effects model showed a main effect of sex (χ^2^_(1)_= 25.17, p<0.001 η^2^=0.08), session (χ^2^_(9)_= 109.04, p<0.001, η^2^=0.03), and a significant interaction sex x session (χ^2^_(9)_=26.86, p<0.01, η^2^=0.007); n=153 males, 162 females. **(D)** Scatter plot showing the relationship between alcohol intake and blood alcohol concentration values. Centerline is the linear regression and dashed lines are the 95% confidence interval. Linear regression F_(1,15)_=7.2, p<0.05, r^2^= 0.36; n=15 mice. **(E)** Diagram depicting the timing of behavioral tests following adolescent alcohol exposure. Animals underwent the DID paradigm from P29 to P40. They were tested 72 hours after the last drinking session (short-term), or after forty days of abstinence (long-term).

### Blood Alcohol Concentration Measurement

Blood alcohol concentration (BAC) was measured in trunk blood immediately after the last drinking session (P40) by using the NAD+/NADH spectrophotometric method, as previously described ^24^.

### Behavioral tests

Mice were handled twice a day for 2 minutes for one week before behavioral test. Mice were placed in the testing room one hour before the beginning of each experimental procedure. All tests were monitored by a camera and analyzed by a blinded experimenter. Apparatus were cleaned with 75% ethanol and dried between mice and sessions. Raw data and movies can be found in Mendeley dataset (DOI: 10.17632/gtnmrbtmt4.1)

### Open field test

Locomotion and anxiety-like behavior were evaluated by conducting the Open Field (OF) test as described in ^25^. Mice were placed into the 40×40×40cm OF and were allowed to explore it freely for 5 minutes (thymotagsis) or 30 minutes (locomotion).

### Elevated Plus Maze Test

Elevated plus maze (EPM) test was conducted as described in ^26^. Animals were placed in the middle of the EPM and were allowed to explore the maze during 5 minutes.

### Forced swimming Test

Forced swimming test was performed according to the procedures described in ^27^. Mice were forced to swim for 6 minutes in a glass cylinder filled with water, and immobility time was recorded during the last 4 minutes of the test.

### Novel Object Recognition Test

NOR was performed as described in ^28^, with a long habituation phase. Mice underwent 3 days of habituation (10 minutes per day), followed by one session of familiarization on day 4, where mice were allowed to explore freely two copies of the same object for 10 minutes. On day 5, one copy of the familiar object was replaced by a novel object and mice were allowed to freely explore their environment for 10 minutes.

### Three Chamber Test

The three chamber test was performed as described in ^29^.

### Reversal Learning Test

The reversal learning test was performed by using the Barnes maze, as described in ^30^, with slight modifications. Learning was assessed for 5 days (2 sessions per day, inter-trial interval one hour). Seventy-two hours after the last learning session, mice underwent the 80-second learning probe trial, in which the escape tunnel was removed from the apparatus. Twenty-four hours after the probe test, mice underwent 5 days (2 sessions per day) of reversal learning. Seventy-two hours after the last reversal learning session, mice underwent the reversal learning probe test.

### Two-bottle choice drinking paradigms

Intermittent access to 20% alcohol (IA20%-2BC) or 1% sucrose (IA1%suc-2BC) two-bottle choice drinking procedures are described in ^23,31^ (Fig 5A).

### Immunohistochemistry

Immunohistochemistry was conducted as previously described ^32^. Fifty μm brain sections were incubated in goat anti-Iba1 antibody overnight at 4C (1/500, Abcam (Cambridge, UK) #ab5076). Donkey anti-goat AlexaFluor 564 was used as secondary antibody. Images were acquired with Nikon A1 confocal microscope. Microglial cell activation state was defined according to their shape: microglia presenting a small nucleus and numerous and ramified extensions were considered as resting, whereas amoeboid-shaped microglial cells with a large nucleus and smaller extensions were considered as activated.

### Western blot analysis

Western blot analysis was conducted from prefrontal cortex extracts as previously described ^32^. Membranes were probed with primary antibodies (rabbit anti-HMGB1 1/1000, abcam #ab18256; rabbit anti-TLR4 1/500, Proteintech (Rosemont, IL, USA) #19811; mouse anti-actin 1/5000, Sigma-Aldrich (Saint Louis, MO, USA) #A3854) overnight at 4C, then probed with HRP-conjugated secondary antibody for one hour at room temperature. Membranes were developed using ECL and images were obtained with ImageQuant LAS 4000 camera system (GE Healthcare, Chicago, IL, USA). Band intensities were quantified using ImageJ software (NIH).

### ELISA

IL-1β levels were determined from prefrontal cortex extracts by using the IL-1β ELISA kit (ThermoFisher Scientific, #88-7013) following the manufacturer’s protocols.

### Statistical tests

Data were analyzed by using two-ways analysis of variance (ANOVA), linear mixed-effects model or student t-test, as detailed in the figure legends. Significant main effects of ANOVA and linear mixed-effects model were further investigated with Tukey *post-hoc* test and statistical significance was set at p<0.05. For results homogeneity purpose, all effect sizes were converted in η^2^ and indicated in figure legends. The number of subjects is indicated in each of the figure legend.

## Results

### Adolescent mice voluntarily binge-drink alcohol and reach high blood alcohol concentrations

Adolescent mice underwent a modified version of the Drinking In the Dark (DID) paradigm ^33^, and were given access to a bottle of alcohol 20% for 4 hours per day, for 10 sessions between P29 and P40 (Fig. 1A). Alcohol intake did not alter body weight gain (Fig. 1B) and adolescent mice voluntarily consumed high amounts of alcohol (Fig. 1C). Our results further showed that females significantly drank more alcohol than males (mean alcohol consumption 7.25±0.2 and 6.52±0.09 g/kg/4h respectively) and exhibited escalation of alcohol consumption (Fig.1C). Blood alcohol concentration (BAC), measured immediately after the last drinking session, positively correlated with alcohol intake (Fig. 1D). Overall, we showed that adolescent mice voluntarily consumed high amounts of alcohol and reached BAC values comprised between 100 and 200mg/dL, which correspond to binge-drinking values observed in humans ^33^. Following AAE, a battery of behavioral tests was performed at two different time-points on independent cohorts of animals: Short-term effects were evaluated 72 hours after the last drinking session, whereas long-term effects were evaluated on adult animals after 40 days of abstinence (Fig. 1E).

### Voluntary adolescent alcohol binge-drinking leads to long-term development of anxiety-like and depressive-like behaviors

Studies have suggested that heavy alcohol exposure in rats leads to the development of anxiety-like behaviors ^34–36^. Following AAE, mice were tested in the Open Field (OF) and the Elevated Plus Maze (EPM) apparatus, which are commonly used to assess anxiety-like behaviors in rodents ^26^ (Fig. 2A, B). Seventy-two hours after the last drinking session, no significant difference in the percentage of time spent in the center of the OF was found between AAE and water-exposed animals over three sessions (Fig 2C). In addition, AAE animals and water littermates spent similar percentage of time exploring the open arm of the EPM (Fig. 2D), and exhibited similar percentage of open arms entries (Fig. 2E).

**Fig 2:**
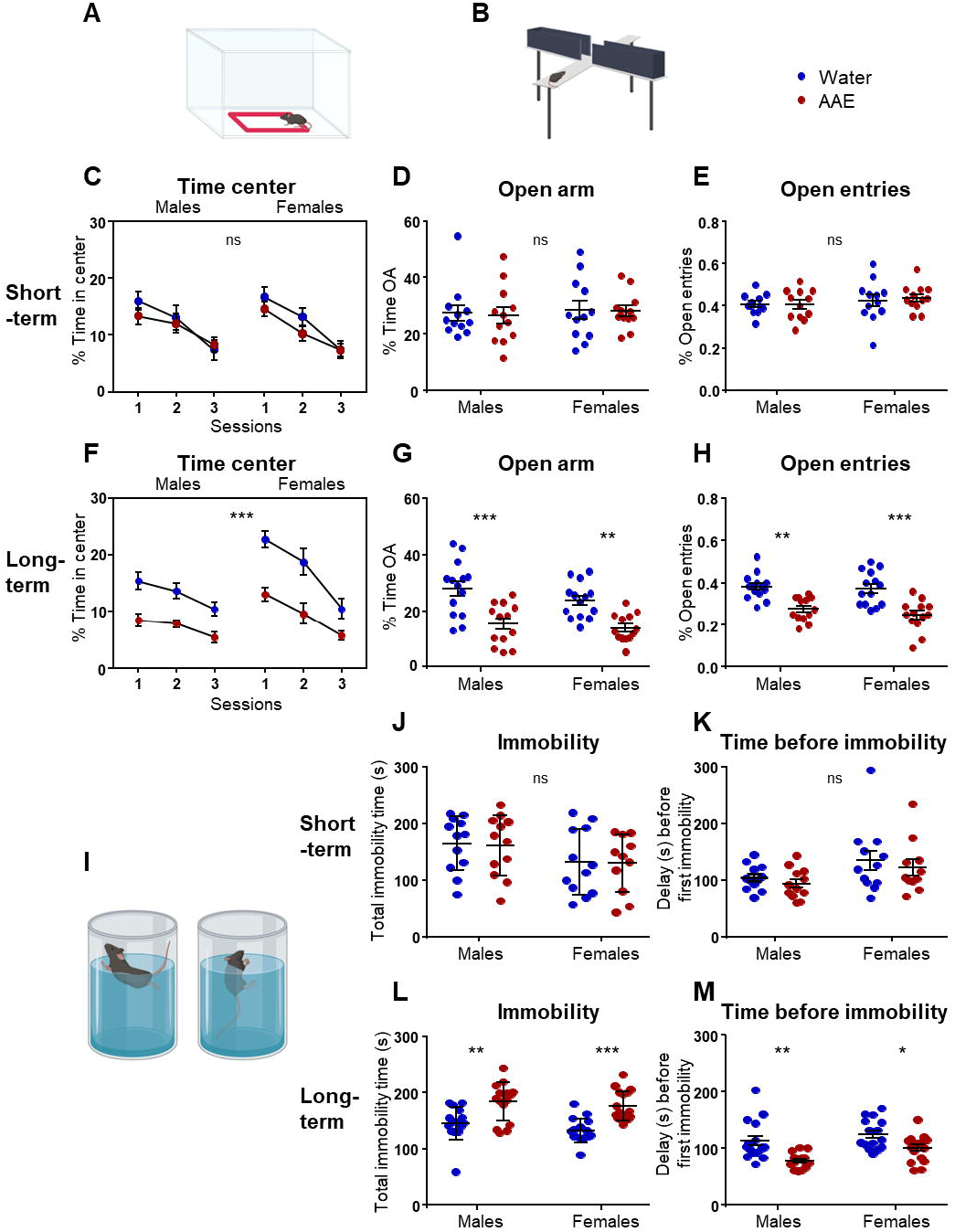
Enhanced anxiety-like and depressive-like behaviors in adulthood but not in adolescence after adolescent alcohol exposure. **(A,B)** Schematic representation of the Open Field (OF) (**A**) and the Elevated Plus Maze (EPM) apparatus (**B**). **(C)** Short-term; Percentage of time spent in the center of the OF. Linear mixed-effects model showed a significant main effect of sessions (χ^2^_(2)_=53.59, p<0.001, η^2^=0.33), but no effect of treatment (χ^2^_(1)_=1.70, p=0.19) or sex (χ^2^_(1)_=0.00, p=0.98) and no interaction; n=12-14 per group. **(D)**Short-term; Percentage of time spent in the open arm of the EPM. Two-way ANOVA showed no main effect of treatment (F_(1,46)_=0.14, p=0.71), or sex (F_(1,46)_=0.66, p=0.42). **(E)** Short-term; Percentage of open arm entries. Two-way ANOVA showed no main effect of Treatment (F_(1,46)_=0.01, p=0.94) or sex (F_(1,46)_=2.52, p=0.12); n=12-14 per group. **(F)** Long-term; Percentage of time spent in the center of the OF. Linear mixed-effects model showed a significant main effect of treatment (χ^2^_(1)_=63.77, p<0.001, η^2^=0.57), sex (χ^2^_(1)_=13.73, p<0.001, η^2^=0.22) and sessions (χ^2^_(2)_=59.09, p<0.001, η^2^=0.44), and a significant interaction sex x session (χ^2^_(2)_=10.08, p<0.01, η^2^=0.07); n=12-14 per group. **(G)** Long-term; Percentage of time spent in the open arm of the EPM. Two-way ANOVA showed a main effect of treatment (F_(1,50)_=30.97, p<0.001, η^2^=0.38), but no main effect of sex (F_(1,50)_=0.89, p=0.35) and no interaction (F_(1,50)_=0.09, p=0.77). *Post-hoc* Tukey test detected a significant difference between Water and AAE in males (p<0.001) and in females (p<0.01). **(H)** Long-term; Percentage of open arm entries. Two-way ANOVA showed a main effect of treatment (F_(1,50)_=40.2, p<0.001, η^2^=0.44), but no main effect of sex (F_(1,50)_=1.10, p=0.30) and no interaction (F_(1,50)_=0.39, p=0.54). *Post-hoc* Tukey test detected a significant difference between Water and AAE in males (p<0.01) and in females (p<0.001); n=12-14 per group. **(I)** Schematic representation of the Forced Swimming Test (FST). Episodes of active swimming (left) and immobility (right) were recorded and analyzed. **(J)** Short-term; Total immobility time. Two-way ANOVA showed a main effect of sex (F_(1,43)_=4.36, p<0.05, η^2^=0.09) but no effect of treatment (F_(1,43)_=0.03, p=0.87) and no interaction (F_(1,43)_=0.003, p=0.96). **(K)** Short-term; Delay before first immobility. Two-way ANOVA showed a main effect of sex (F_(1,43)_=6.17, p<0.05; η^2^=0.12) but no effect of treatment (F_(1,43)_=1.05, p=0.31) and no interaction (F_(1,43)_=0.01, p=0.94); n=11-12 per group. **(L)** Long-term; Total immobility time. Two-way ANOVA showed a main effect of treatment (F_(1,61)_=35.21, p<0.001, η^2^=0.36) but no effect of sex (F_(1,61)_=2.39, p=0.13) and no interaction (F_(1,61)_=0.11, p=0.74). *Post-hoc* Tukey test revealed a significant difference between AAE and water mice, both in males (p<0.01) and in females (p<0.001). **(M)** Long-term; Delay before first immobility. Two-way ANOVA showed a main effect of treatment (F_(1,61)_=22.98, p<0.001, η^2^=0.25) and sex (F_(1,61)_=7.77, p=0.007, η^2^=0.08) but no interaction (F_(1,61)_=0.66, p=0.42). *Post-hoc* Tukey test revealed a significant difference between AAE and water mice, both in males (p<0.01) and in females (p<0.05); n=16-17 per group.

Anxiety-like behavior was further assessed on independent cohorts of mice after 40 days of abstinence. Our data revealed that adult abstinent mice that were exposed to alcohol during adolescence exhibited significantly enhanced thymotagsis in the OF as compared to water littermates, in both sexes (Fig. 2F). Indeed, despite exhibiting habituation across the 3 consecutive sessions, AAE animals spent significantly less time than the water controls in the center of the OF (Fig. 2F). In addition, AAE animals spent less time exploring the open arms of the EPM (Fig. 2G), and the percentage of open arm entries was reduced as compared to water animals (Fig. 2H). Importantly, no significant difference in locomotor activity or habituation was found between AAE and water mice (Fig. S1A, C), and the total number of EPM arm entries did not differ between groups, suggesting that AAE did not impact exploration behavior (Fig. S1B, D). Together, these data suggest that although voluntary binge-drinking of alcohol during adolescence did not alter anxiety levels in adolescence, it promoted the long-term development of anxiety-like behaviors.

We then assessed the consequences of AAE on depressive-like behavior by using the Forced Swimming Test (FST), which is widely used to investigate the response to antidepressant treatments and assess depressive-like behavior in animal models ^27^ (Fig. 2I). Short-term after the last drinking session, AAE and water adolescent mice exhibited equivalent immobility time and delay before the first immobility, both in males and in females (Fig. 2J, K). Interestingly, adolescent males exhibited higher immobility time and decreased delay before first immobility, as compared to females (Fig. 2J, K).

In contrast, when the FST was performed 40 days after the last alcohol drinking session, mice exposed to AAE exhibited significantly increased immobility time compared to water littermates, in both sexes (Fig. 2L). In addition, the delay before the first immobility episode was shorter in AAE animals (Fig. 2M), suggesting that AAE induces the development of depressive-like behavior long-term after alcohol consumption.

### Decreased novel object exploration in adult mice exposed to alcohol binge-drinking during adolescence

The Novel Object Recognition (NOR) test assesses the natural preference for novel objects normally displayed by mice, and gives insights about their recognition memory performance (Fig. 3A) ^37^. Shortly after AAE, all groups of adolescent mice showed similar exploration behavior (Fig. 3B, C). In addition, no significant difference in discrimination index (DI) and familiar object habituation index (FHI) was found between animals (Fig. 3D, E).

**Fig 3:**
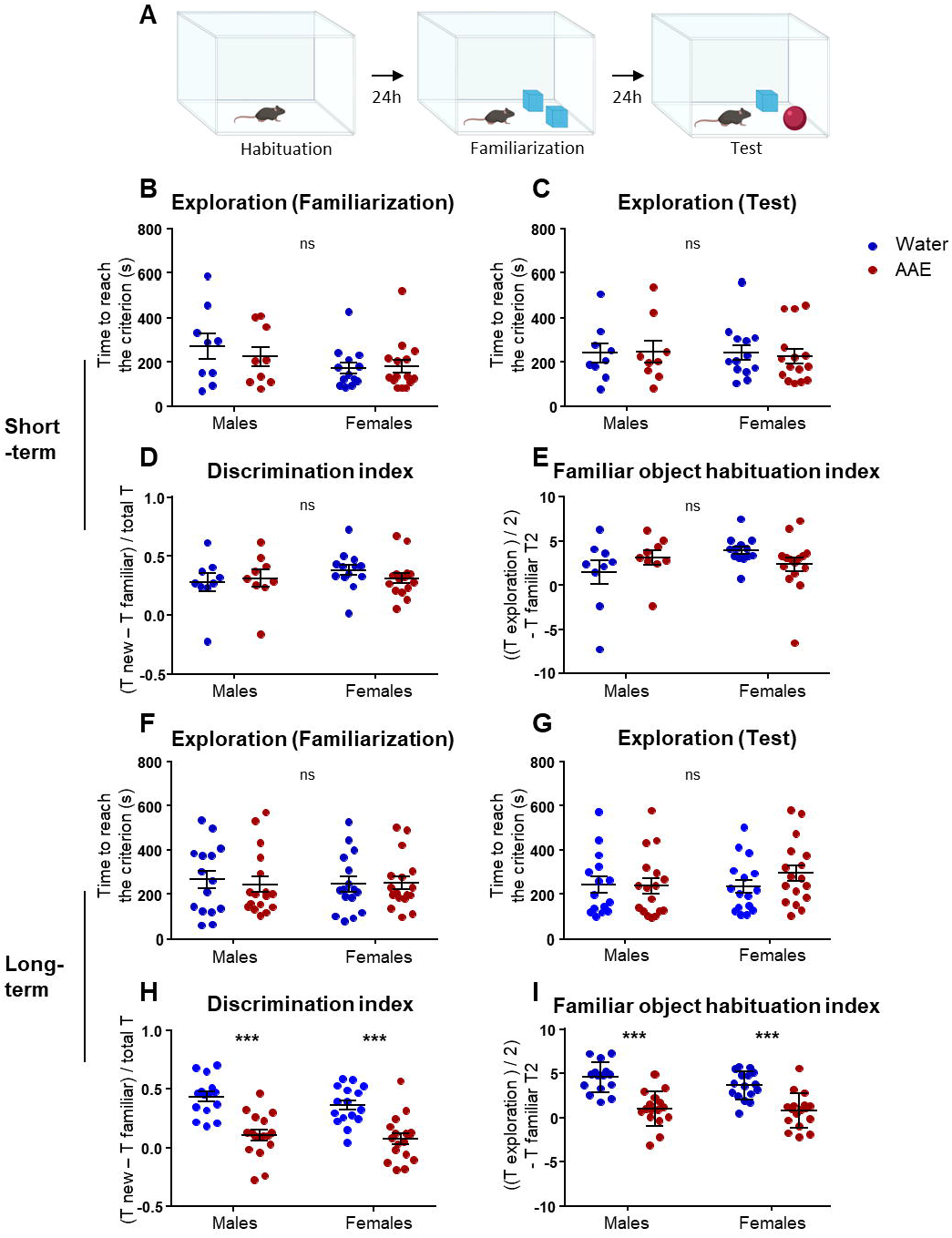
Adolescent binge-drinking decreases novel object recognition performances in adult but not adolescent mice. **(A)** Schematic representation of the Novel Object Recognition test (NOR). Following 3 days of habituation in the open field, mice were allowed to familiarize with two copies of the same object for 10 minutes. Twenty-four hours later, one copy of the familiar object is replaced by a novel object and exploration time is recorded. **(B)** Short-term; Familiarization session: Time to reach criterion (20 seconds of total object exploration). Two-way ANOVA showed no main effect of treatment (F_(1,42)_=0.27, p=0.6) or sex (F_(1,42)_=3.52, p=0.07). **(C)** Short-term; Test session: Time to reach criterion. Two-way ANOVA showed no main effect of treatment (F_(1,42)_=0.02, p=0.88) or sex (F_(1,42)_=0.05, p=0.83). **(D)** Short-term; Discrimination Index (DI), calculated as the time exploring the novel object minus the time exploring the familiar object, divided by the total exploration time (~20 seconds). Two-way ANOVA showed no main effect of treatment (F_(1,42)_=0.10, p=0.76) or sex (F_(1,42)_=0.87, p=0.36). **(E)** Short-term; Index of habituation to the familiar object (FHI), calculated as the time exploring both objects during familiarization/2, minus time exploring familiar object during test session. Two-way ANOVA showed no main effect of treatment (F_(1,42)_=0.12, p=0.73) or sex (F_(1,42)_=1.06, p=0.31); n=9-15 per group. **(F)** Long-term; Familiarization session: Time to reach criterion (20 seconds of total object exploration). Two-way ANOVA showed no main effect of treatment (F_(1,60)_=0.03, p=0.87) or sex (F_(1,60)_=0.1, p=0.76). **(G)** Long-term; Test session: Time to reach criterion. Two-way ANOVA showed no main effect of treatment (F_(1,60)_=0.81, p=0.37) or sex (F_(1,60)_=0.39, p=0.53). **(H)** Long-term; Discrimination index. Two-way ANOVA showed a significant main effect of treatment (F_(1,60)_=49.86, p<0.001, η^2^=0.45) but not sex (F_(1,60)_=1.15, p=0.29) and no interaction (F_(1,60)_=0.33, p=0.57). *Post-hoc* Tukey test revealed a significant difference between AAE and water-exposed animals in both males and females (p<0.001). **(I)** Long-term; Index of habituation to the familiar object (HFI). Two-way ANOVA showed a significant main effect of treatment (F_(1,60)_=49.50, p<0.001, η^2^=0.44) but not sex (F_(1,60)_=1.61, p=0.21) and no interaction (F_(1,60)_=0.55, p=0.46). *Post-hoc* Tukey test revealed a significant difference between AAE and water-exposed animals, in both males and females (p<0.001); n=15-17 per group.

However, 40 days after the last alcohol exposure, despite similar exploration behavior, AAE mice exhibited significantly lower DI and FHI compared to water littermates, both in males and females (Fig. 3F-I). Those findings suggest that although AAE did not impact recognition memory performance in adolescence, it dramatically impaired novel object recognition in adulthood.

### Adolescent alcohol binge-drinking does not affect mouse sociability

We investigated consequences of AAE on social behaviors by performing the three-chambered social approach task ^29^ (Fig. S2A). All mice showed similar preference for the social stimulus, suggesting that AAE did not impact mice sociability (Fig. S2B, D). Social novelty preference was also investigated by calculating the social novelty preference index (SNI) after introduction of a stranger mouse in the empty wired cup (Fig. S2A), but no significant difference in social novelty preference between AAE and water adolescent mice was observed, either short- or long-term after AAE (Fig. S2C, E). However, in contrast to adolescents, adult mice failed to exhibit significant social novelty preference (SNI not significantly different from zero). Overall, those results suggest that AAE had no major impact on mouse sociability.

### Impaired reversal learning long-term after adolescent alcohol binge-drinking

Clinical studies have reported that AAE may lead to long-lasting deficits in executive functions ^11,12^. We sought to unveil the consequences of AAE on reversal spatial learning by performing the Barnes Maze test (Fig. 4A). Learning abilities were first investigated. Shortly after the last drinking session, escape time were not different between AAE- and water animals, and similarly decreased over days (Fig. S3A). Accordingly, the number of primary errors also followed a similar decrease over sessions in all groups (Fig. S3B). Three days later, a probe test session in absence of the escape tunnel was performed, and AAE and water animals spent similar percentage of time spent in the correct sector of the maze (Fig. S3C). Long-term after AAE, similarly to adolescent mice, learning abilities of adult mice were not affected by AAE, and similar decreasing escape time and primary errors were observed in all groups (Fig. S3D, E). Moreover, no difference was found during the probe test (Fig. S3F). Together, these results suggest that AAE had no effect on spatial learning acquisition.

**Fig 4:**
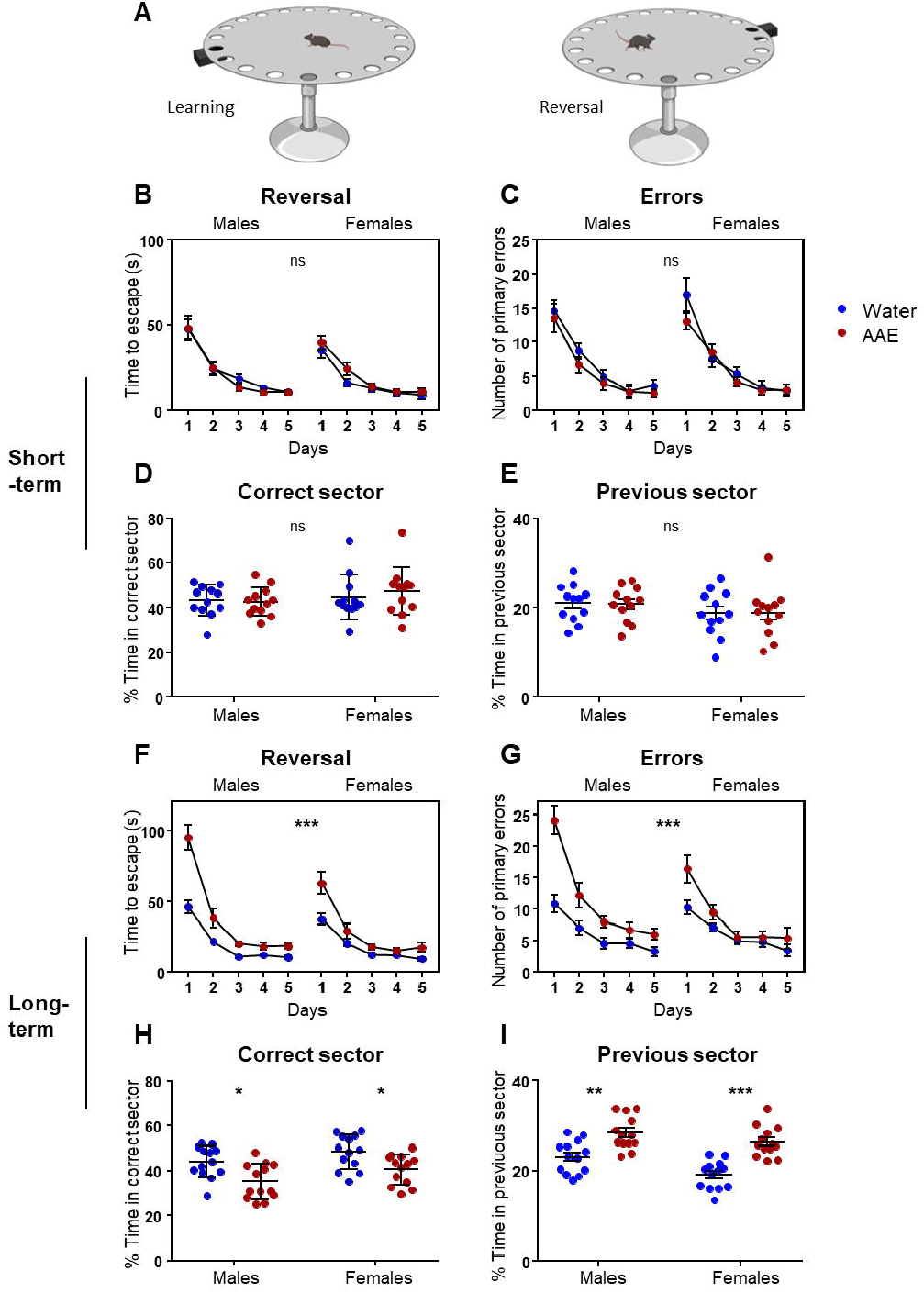
Behavioral flexibility is impaired long-term after adolescent alcohol exposure. **(A)** Schematic representation of the Barnes maze test. During learning, mice are given 10 training sessions in order to learn the position of the escape tunnel (left, see supplemental figure 3). Then, the position of the escape tunnel is modified and mice are given 10 reversal learning sessions (right). **(B)** Short-term; Reversal learning: mean escape time per day across 5 days. Linear mixed-effects model showed a significant main effect of day (χ^2^_(4)_=363.3, p<0.001, η^2^=0.55) and sex (χ^2^_(1)_=4.52, p<0.05, η^2^=0.086) but no main effect of treatment (χ^2^_(1)_=0.26, p=0.61) and no interaction (day x sex χ^2^_(4)_=7.90, p=0.1; day x treatment χ^2^_(4)_=3.37, p=0.5; sex x treatment χ^2^_(1)_=1.64, p=0.2). **(C)** Short-term; Reversal learning: primary errors per day across 5 days. Linear mixed-effects model showed a significant main effect of day (χ^2^_(4)_=342.42, p<0.001, η^2^=0.55), but no main effect of treatment (χ^2^_(1)_=2.15, p=0.14) or sex (χ^2^_(1)_=0.32, p=0.57) and no interaction (day x sex χ^2^_(4)_=0.54, p=0.97; day x treatment χ^2^_(4)_=3.11, p=0.54; sex x treatment χ^2^_(1)_=0.01, p=0.91). **(D)** Short-term; Percentage of time spent in the correct sector during probe test, 72 hours after the last reversal learning session. Two-way ANOVA showed no main effect of treatment (F_(1,44)_=0.18, p=0.67) or sex (F_(1,44)_=1.47, p=0.23). **(E)** Short-term; Percentage of time spent in the previous sector during probe test. Two-way ANOVA showed no main effect of treatment (F_(1,44)_=0.01, p=0.93) or sex (F_(1,44)_=2.49, p=0.12); n=12 per group. **(F)** Long-term; Reversal learning: mean escape time per day. Linear mixed-effects model showed a significant main effect of day (χ^2^_(4)_=539.37, p<0.001, η^2^=0.64), treatment (χ^2^_(1)_=47.87, p<0.001, η^2^=0.22) and sex (χ^2^_(1)_=7.92, p<0.001, η^2^=0.13), as well as a significant interaction day x sex (χ^2^_(4)_=24.81, p<0.001, η^2^=0.08), and day x treatment (χ^2^_(4)_=61.63, p<0.001, η^2^=0.17) but not treatment x sex (χ^2^_(1)_=3.42, p=0.06). **(G)** Long-term; Reversal learning: primary error number. Linear mixed-effects model showed a significant main effect of day (χ^2^_(4)_=273.91, p<0.001, η^2^=0.46), treatment (χ^2^_(1)_=29.15, p<0.001, η^2^=0.16) and sex (χ^2^_(1)_=4.10, p<0.05, η^2^=0.07), as well as an interaction day x treatment (χ^2^_(4)_=39.74, p<0.001, η^2^=0.11), sex x treatment (χ^2^_(1)_=4.51, p<0.05, η^2^=0.03) but not day x sex (χ^2^_(4)_=8.78, p=0.07). **(H)** Long-term; Percentage of time spent in the correct sector during probe test, 72 hours after the last reversal learning session. Two-way ANOVA showed a significant main effect of treatment (F_(1,48)_=16.62, p<0.001, η^2^=0.24) and sex (F_(1,48)_=6.01, p<0.05, η^2^=0.09) but no interaction (F_(1,48)_=0.07, p=0.79). *Post-hoc* Tukey test revealed a significant difference between AAE and water animals in males (p<0.05) and in females (p<0.05). **(I)** Long-term; Percentage of time spent in the previous sector during probe test. Two-way ANOVA showed a significant main effect of treatment (F_(1,48)_=44.19, p<0.001, η^2^=0.43) and sex (F_(1,48)_=9.91, p<0.01, η^2^=0.1) but no interaction (F_(1,48)_=1.19, p=0.28). *Post-hoc* Tukey test revealed a significant difference between AAE and water animals in males (p<0.01) and in females (p<0.001); n=13 per group.

We next assessed reversal learning abilities by rotating the escape tunnel location by 180° (Fig. 4A). Adolescent AAE and water mice displayed similar escape time, and no difference was found in the number of primary errors (Fig. 4B, C). Furthermore, the probe test did not reveal any difference in the percentage of time spent in the correct sector (Fig. 4D) or in the previous sector (Fig. 4E).

However, when reversal learning was assessed in adulthood, AAE-exposed mice showed higher escape time and made more errors as compared to water controls, in both sexes (Fig. 4F, G). Interestingly, the mean escape time and primary error number were also significantly higher in males compared to females, regardless of their alcohol treatment (Fig. 4F, G). In addition, during the probe test, AAE mice spent significantly less time in the correct sector compared to water controls (Water males 44.04±1.94%, AAE males 35.24±2.18%, Water females 48.44±2.09%, AAE females 40.75±1.86%) (Fig. 4H) and more time in the sector corresponding to the previous position of the escape tunnel (Water males 23.16±3.6%, AAE males 28.4±3.6%, Water females 19.15±3.1%, AAE females 26.46±3.3%), suggesting increased perseveration behavior in AAE groups (Fig. 4I). Altogether, our results suggest that while AAE did not alter reversal learning short-term after the last drinking session, it strongly impaired it in adult mice, long-term after alcohol exposure.

### Adolescent alcohol binge-drinking enhances alcohol consumption in adulthood

Clinical and pre-clinical studies have suggested that AAE increases the risk of developing alcohol use disorders later in life ^5,10^. In order to decipher whether AAE modulates alcohol intake and preference, mice underwent 5 sessions of intermittent access to 20% alcohol – 2 bottle choice paradigm (Fig. 5A) ^31,32^. Alcohol intake, alcohol preference and total fluid intake were measured at the end of each 24-hour session. Short-term after AAE, no significant difference was observed in alcohol intake or preference between groups (Fig. 5B-D). Interestingly, data revealed that adolescent females tend to consume more alcohol than males (Mean alcohol intake 26.92±2.96 and 23.95±2.03 g/kg/24h respectively), and more fluid in total (Mean fluid intake 310.5±6.12 and 265.9±24.5 ml/kg/24h, respectively) (Fig. 5B-D).

**Fig 5:**
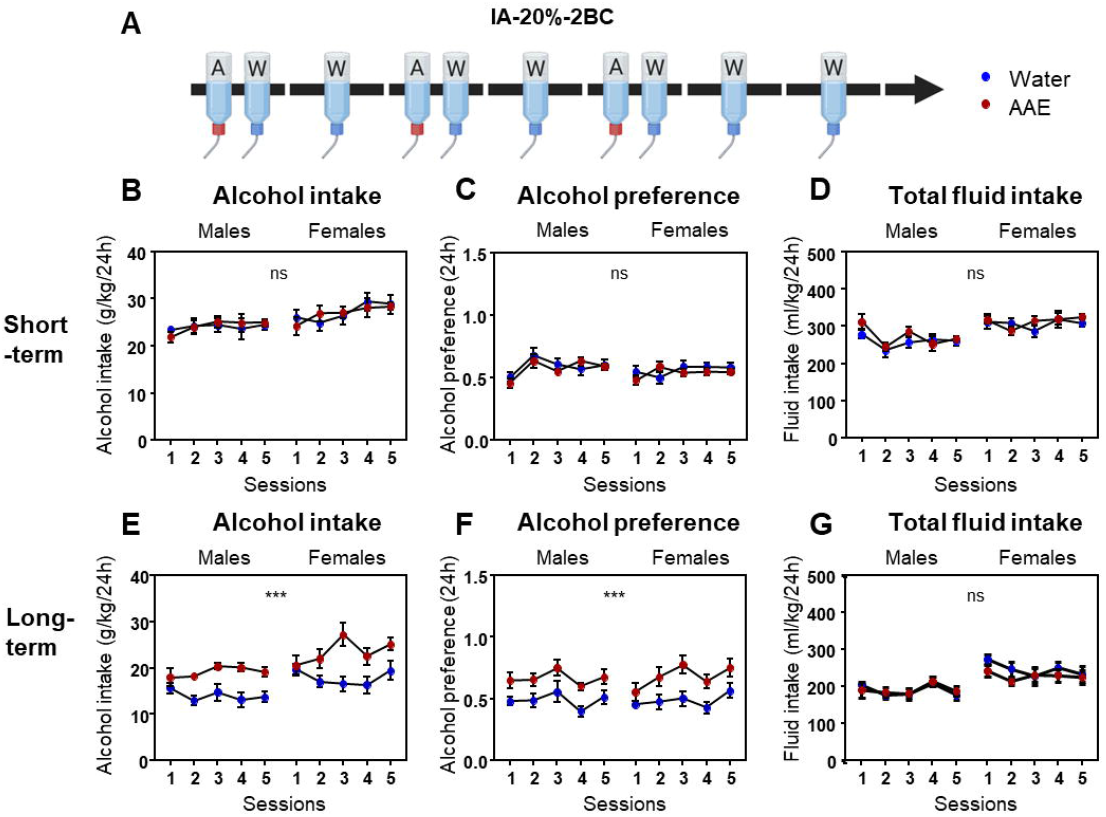
Binge-drinking during adolescence increases alcohol consumption and preference in adult mice. **(A)** Scheme depicting the intermittent access to alcohol 20%-two bottle choice paradigm. Short-term (B-D) and Long-term (E-G) after AAE, mice underwent the IA-20%-2BC paradigm for 5 sessions. **(B)** Short-term; Alcohol intake (g/kg/24h). Linear mixed-effects model showed a significant main effect of session (χ^2^_(4)_=11.3, p<0.05, η^2^=0.04), and sex (χ^2^_(1)_=17.28, p<0.001, η^2^=0.27) but no effect of treatment (χ^2^_(1)_=0.1, p=0.76) and no interaction (session x treatment χ^2^_(4)_=1.4, p=0.85; sex x treatment χ^2^_(1)_=0.08, p=0.78; session x sex χ^2^_(4)_=2.88, p=0.58). **(C)** Short-term; Alcohol preference. Linear mixed-effects model showed a significant main effect of session (χ^2^_(4)_=22.42, p<0.001, η^2^=0.07), but no effect of sex (χ^2^_(1)_=1.86, p=0.17) or treatment (χ^2^_(1)_=0.85, p=0.36) and no interaction (session x treatment χ^2^_(4)_=4.57, p=0.33, sex x treatment χ^2^_(1)_=0.00, p=0.96, session x sex χ^2^_(4)_=9.35, p=0.051). **(D)** Short-term; Total fluid intake (ml/kg/24h). Linear mixed-effects model showed a significant main effect of session (χ^2^_(4)_=15.12, p<0.01, η^2^=0.05) and sex (χ^2^_(1)_=34.6, p<0.001, η^2^=0.42) but no effect of treatment (χ^2^_(1)_=1,78 p=0.18) and no interaction (session x treatment χ^2^_(4)_=5.53, p=0.24, sex x treatment χ^2^_(1)_=0.27, p=0.6, session x sex χ^2^_(4)_=7.78, p=0.1); n=12 per group. **(E**) Long-term; Alcohol intake (g/kg/24h). Linear mixed-effects model showed a significant main effect of treatment (χ^2^_(1)_=34.86, p<0.001, η^2^=0.15) and sex (χ^2^_(1)_=17.28, p<0.001, η^2^=0.09) but no effect of session (χ^2^_(4)_=8.53 p=0.07). This model also showed an interaction session x treatment (χ^2^_(4)_=14.8, p=0.01, η^2^=0.02) but not sex x treatment (χ^2^_(1)_=0.00, p=0.98) or session x sex χ^2^_(4)_=3.37, p=0.50). **(F)** Long-term; Alcohol preference. Linear mixed-effects model showed a significant main effect of treatment (χ^2^_(1)_=33.21, p<0.001, η^2^=0.15) and session (χ^2^_(4)_=19.6, p<0.001, η^2^=0.06) but no effect of sex (χ^2^_(1)_=0.04 p=0.84) and no interaction (session x treatment χ^2^_(4)_=2.17, p=0.7, sex x treatment χ^2^_(1)_=0.01, p=0.92, session x sex χ^2^_(4)_=3.32, p=0.51). **(G)** Long-term; Total fluid intake (ml/kg/24h). Linear mixed-effects model showed a significant main effect of session (χ^2^_(4)_=14.63, p<0.01, η^2^=0.04) and sex (χ^2^_(1)_=21.1, p<0.001, η^2^=0.1) but no effect of treatment (χ^2^_(1)_=0.3 p=0.58) and no interaction (session x treatment χ^2^_(4)_=2.69, p=0.61, sex x treatment χ^2^_(1)_=1.04, p=0.31, session x sex χ^2^_(4)_=3.39, p=0.49); n=12 per group.

In contrast, adult mice which were exposed to alcohol during adolescence consumed significantly more alcohol than water littermates in both sexes (Mean alcohol intake: Water males 14.0±2.7; AAE males 19.12±1.54; Water females 17.78±3.2; AAE females 23.45±5.3 g/kg/24h) (Fig. 5E). AAE mice also exhibited higher alcohol preference (Fig. 5F), without any difference in total fluid intake (Fig. 5G). Similarly, females significantly consumed higher amount of alcohol compared to males (Mean alcohol consumption 20.61±5.2 and 16.56±3.4 g/kg/24h, respectively) and presented higher total fluid intake (Mean fluid intake 236.6±43 and 189.7±27 ml/kg/24h, respectively) (Fig. 5E-G).

Furthermore, sucrose consumption was measured short- and long-term after AAE, and no difference in sucrose intake or preference was found between AAE animals and water controls in both sexes (Fig S4).

### Voluntary alcohol binge-drinking during adolescence does not induce major induction of neuro-inflammation in the prefrontal cortex

Several studies using binge-like administration of alcohol during adolescence reported alcohol-induced activation of innate immune signaling in the frontal cortex and promoted behavioral alterations similar to those observed in this study ^18–20,38^. In order to decipher whether voluntary alcohol binge-drinking in adolescent mice induces neuro-inflammation in the PFC, we assessed the number and activation state of microglial cells by immunofluorescence. Microglial cells with small nucleus and numerous ramified extensions were considered as resting, whereas amoeboid-shaped microglial cells with a large nucleus and smaller extensions were considered as activated ^39^ (Figure S5A, B). As shown in Figure 6A-E, there was no difference in the number of activated microglial cells in the prelimbic cortex between AAE and water animals when analyzed 72 hours after the least drinking session (Figure 6E), and no difference was found in the total number of microglial cells (Figure S5C). The same analysis was performed in the PFC of adult animals and no alcohol-dependent activation of microglia was observed (Figure S5D-I). Moreover, no difference was observed in the infralimbic region of the PFC (data not shown). As passive administration of alcohol in adolescent rodents has been shown to increase the expression of the High Mobility Group protein B1 (HMGB1) and the Toll-like receptor 4 (TLR4) in the PFC ^40,41^, we further assessed their expression after voluntary alcohol binge-drinking. Surprisingly, as shown in figure 6F and 6G, no significant increase in HMGB1 or TLR4 expression was observed in the PFC of AAE animals compared to water controls, neither in males or females, short-term or long-term after AAE. Finally, we assessed the concentration of interleukin-1β in the PFC of AAE and water mice and did not find any difference between groups, neither 3 days nor 40 days after AAE (Figure 6H, I). Altogether, our data suggest that our model of voluntary binge-drinking of alcohol in mice did not induce major neuro-inflammation in the PFC.

**Fig 6:**
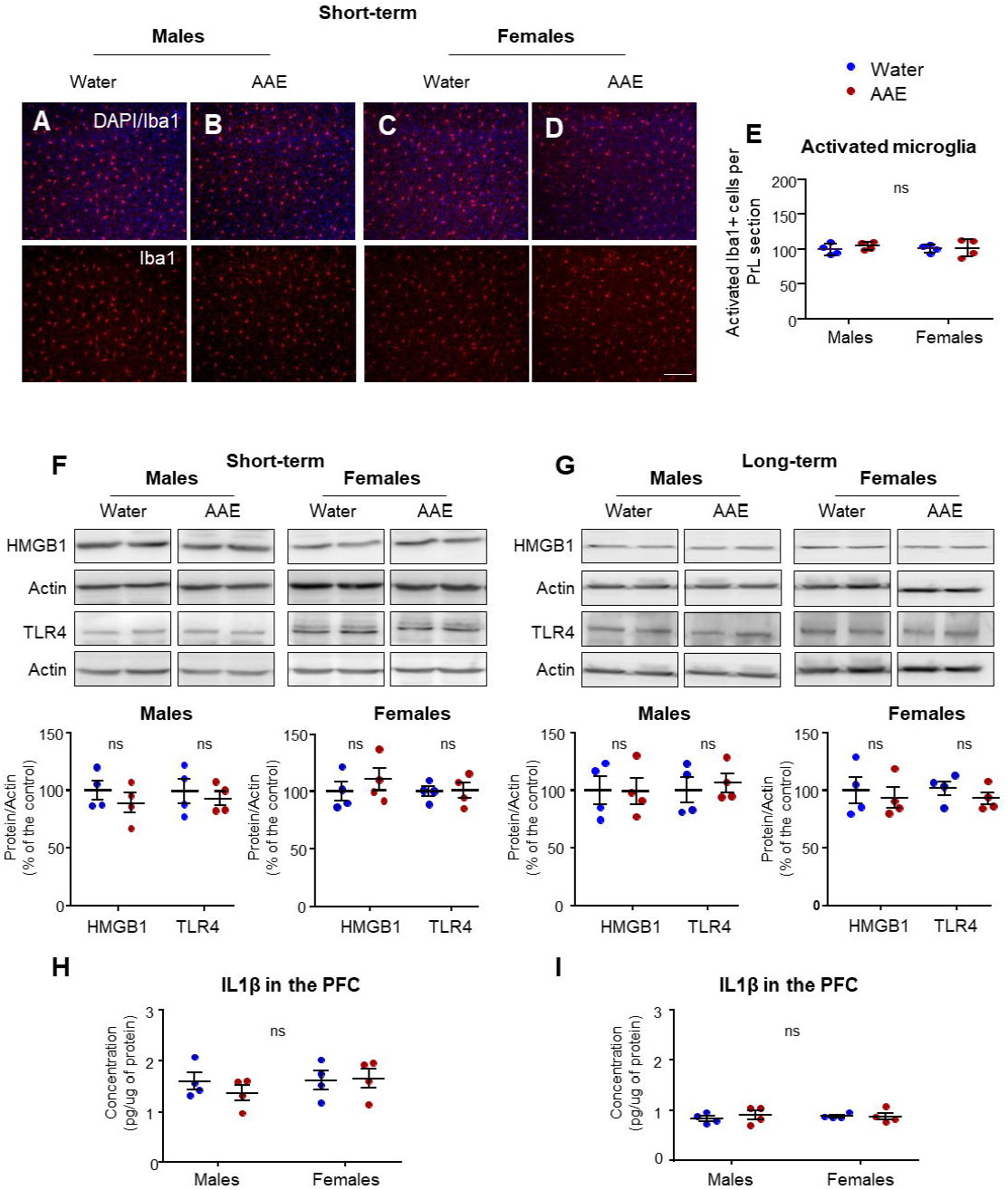
Voluntary alcohol binge-drinking during adolescence does not trigger innate immune system activation in the prefrontal cortex. **(A-D)** Short-term; morphological analysis of microglial cells expressing Iba1 (red) in the prelimbic prefrontal cortex of males water (**A**), males AAE (**B**), females water (**C**) and females AAE (**D**) by immunofluorescence; Bar scale 10μm. **(E)** Short-term; Number of activated Iba1+ microglial cells per prelimbic PFC section (mean ± S.E.M.; Cell body to cell size ratio between 12 and 15). Two-way ANOVA showed no main effect of treatment (F_(1,12)_=0.47, p=0.5), sex (F_(1,12)_=0.05, p=0.82) and no interaction (F_(1,12)_=0.22, p=0.65). n=4 per group. **(F, G)**HMGB1 and TLR4 protein expression were determined by western blot analysis short-term (**F**) and long-term (**G**) after adolescent alcohol exposure. ImageJ was used for optical density quantification. Data are expressed as the average ratio ± S.E.M. of HMGB1 or TLR4 to actin and are expressed as percentage of water control. Significance was determined using two-tailed unpaired t-test. n=4 per group. **(H, I)** IL-1β concentration (pg/μg of protein) in the PFC, short-term (**H**) or long-term (**I**) after adolescent alcohol exposure. **(H)** Short-term; Data are represented as the mean concentration ± S.E.M. Two-way ANOVA showed no main effect of treatment (F_(1,12)_=0.29, p=0.6), sex (F_(1,12)_=0.78, p=0.39) and no interaction (F_(1,12)_=0.61, p=0.45). **(I)** Long-term; Data are represented as the mean concentration ± S.E.M. Two-way ANOVA showed no main effect of treatment (F_(1,12)_=0.25, p=0.62), sex (F_(1,12)_=0.03, p=0.87) and no interaction (F_(1,12)_=0.35, p=0.57). n=4 per group.

## Discussion

### AAE induces anxiety and depressive-like behaviors in adult mice

We report that voluntary adolescent alcohol binge-drinking promotes the development of anxiety- and depressive-like behaviors in adulthood, without any warning sign in adolescence. Such results are consistent with other studies showing that repeated passive exposure to alcohol during adolescence induces the development anxiety-like behaviors in adult rodents ^34–36,42,43^. However, opposite findings have been reported, showing that ethanol vapor exposure during adolescence increased exploration of adult rats in the open arms of the EPM; and such results have been interpreted as AAE-dependent increased impulsivity ^44–46^. OF and EPM are tasks assessing the balance between the innate exploratory drive and the anxiety generated by a novel environment ^26^. Therefore, it remains challenging to decipher whether behavioral difference result from change in anxiety and/or impulsivity, and data should be carefully interpreted. Furthermore, our results also suggest that AAE, while not impacting adolescent behavior in the FST, led to the long-term development of depressive-like behaviors. Such results are in line with a study conducted by Lee *et al*, in which the authors used a similar mouse model and showed AAE-dependent increased depressive-like behaviors and anhedonia in adult males ^47^.

### AAE decreases recognition memory performances in adulthood

The novel object recognition (NOR) test is commonly used for assessing the effects of a drug on memory performance because no reward or reinforcement are needed; as such, the test relies primarily on the rodent’s innate exploratory behavior ^28,37^. As environmental familiarization may modulate novel object interaction, increased anxiety levels in AAE animals could participate in the AAE-dependent impaired recognition memory ^28^. In order to minimize potential bias, we used a long habituation protocol which reduces the stress associated with the OF. Importantly, no difference was found between AAE and water mice regarding their exploratory behavior, and no preference for one object was evidenced. Intriguingly, Pascual *et al* showed reduced performances in the NOR test in both late adolescent and adult rats after AAE ^19^, and they recently reported AAE-induced decreased recognition memory in late adolescent mice ^48^. In opposition, in the present study we reported impaired recognition memory in adult mice only. Such discrepancy may arise from experimental set up differences, including the animal model (mice vs rats), the timing of behavioral testing and the mode of alcohol administration. Indeed, in our model, adolescent mice had access to alcohol between P29 and P40 and were tested at P43, whereas in Pascual *et al*’s study, mice were tested at P46 ^48^. Adolescence is a relatively short developmental period, characterized by a rapid maturation of the frontal brain regions, and is commonly divided into 3 distinct periods: early (PN21-34), middle (PN34-46) and late adolescence/early adulthood (PN46-59) ^22^. It is thus possible that the defects observed in recognition memory at P46 are not yet detectable at P43. Moreover, different administration routes lead to large differences in alcohol pharmacokinetics. Indeed, alcohol is more rapidly absorbed after i.p. as compared to oral ingestion, and alcohol accumulation in the brain is also lower after oral administration ^49^. Such differences must also be taken into account when comparing different results, and it is possible that the timing of appearance of behavioral defects depends on the final alcohol concentration administered to the brain.

### AAE does not lead to sociability defects

Very few studies examined the effects of AAE on sociability. In the present study, we found that AAE does not impair sociability when assessed by the 3-chamber test, neither short- or long-term after AAE. Interestingly, Sabry *et al* showed that AAE did not affect sociability in adolescent male rats, but suppressed sociability when coupled to overcrowding conditions ^50^. This suggests that alcohol exposure *per se* may not be sufficient to affect sociability, but rather enhances the development of social issues when associated with social stress ^50^. We also showed that AAE does not alter the social novelty preference in adolescence. However, in adult mice, as none of the groups of mice exhibited significant social novelty preference, no conclusion can be drawn from our findings. This is probably due to the mouse strain used in our study, as it has been shown that adult C57Bl6 mice lacked to demonstrate social novelty preference in the three-chamber apparatus ^51^.

### AAE impairs behavioral flexibility in adulthood but not in late adolescence

Several cognitive studies have suggested that AAE has minimal effect on spatial learning and memory tasks ^11^. However, flexibility impairments have been reported when reversal learning or set-shifting tasks were demanded ^11,43,46^. Accordingly, we showed that AAE has no effect on spatial learning, but significantly impairs reversal learning for spatial tasks in adulthood. Indeed, all mice were able to learn the initial position of the escape tunnel, but when the task required a more flexible strategy, AAE animals significantly lacked behavioral flexibility and exhibited increased perseveration of previously learned behavior. Such defect in flexibility has been interpreted as a loss of executive functions caused by disruption of frontal cortical areas control ^6,11,38^. Here we show that AAE-dependent impairment of behavioral flexibility is a long-term developmental process, and that frontal brain regions alterations may not be present in late adolescence yet, but rather develop along with frontal circuit maturation.

### AAE enhances adult but not adolescent alcohol consumption and preference

AAE has also been reported to promote alcohol consumption in adult rats ^11,35,38,46,47^, although opposite results have been reported ^52,53^. In the present study, voluntary adolescent binge-drinking promoted alcohol consumption and preference in adult males and females, but not in adolescents. This is quite surprising to see that naïve, water-drinking adolescent mice consumed the same amount of alcohol than animals previously exposed to alcohol, and those results should be carefully interpreted. Indeed, we cannot exclude the possibility that the absence of significant difference between AAE and water-exposed mice might be due to a plateau effect as adolescent C57Bl6 mice voluntarily consume very high amounts of alcohol. Interestingly, sucrose consumption and preference were not affected by AAE, suggesting that the mechanisms triggered by alcohol in the adolescent brain are not shared by all rewarding substances.

### Delayed appearance of behavioral defects: potential mechanisms

This study reports that although voluntary alcohol binge-drinking during adolescence does not lead to short-term behavioral alterations, it dramatically impairs adult executive functions and behavior. This suggests that alcohol perturbs brain maturation to promote the progressive development of behavioral defects, which only emerge during adulthood. It is important to note that all the behaviors assessed in this study involve the PFC, and as such, the impaired behaviors observed during adulthood are likely to arise from PFC malfunction, which may result from alcohol-dependent interference with its maturation ^11,38,54^. Adolescence is a particularly vulnerable period regarding the consequences of alcohol exposure, and the peak of PFC maturation is observed precisely during this developmental period ^2,5^. Although the consequences of AAE on PFC maturation and function are not fully understood yet, mechanistic findings in brain circuitry underlying AAE-induced behavioral impairments are starting to emerge^6,11,38,54^.

Several studies have reported that repeated passive exposure to high levels of alcohol during adolescence induced the long-lasting induction of innate immune signaling through signaling cascades involving RAGE, HMGB1, Toll-like receptors and pro-inflammatory cytokines, which in turn led to synaptic plasticity disruption and neuropathy in frontal brain regions ^19,20,40,42,55^. In addition, in an elegant study, Montesinos *et al* showed that TLR4 KO mice were protected against the alcohol-induced behavioral alterations ^42^. Surprisingly, in the present study, despite severe behavioral defects following AAE, no major induction of neuro-inflammation was observed in the mouse PFC. Experimental differences, such as administration mode of alcohol, may explain discrepancies between studies. Indeed, alcohol absorption profiles appear very different in the two mouse models of AAE: in the voluntary alcohol consumption model, mice drank around 7g/kg of alcohol in a period of 4 hours, whereas i.p. injections involved a single acute administration of 3-5g/kg of alcohol. As elimination of alcohol is done at a constant rate ^56^, the maximal concentration of alcohol after i.p injection is significantly higher as compared to voluntary drinking. It is thus possible that alcohol reaches a toxicity threshold and triggers the induction of neuro-inflammation in frontal brain regions, which is not observed after voluntary alcohol drinking. In addition, it is important to question the role of stress, which is likely to be associated with gavage or i.p. administration of alcohol. Indeed, multiple interactions between stress and alcohol have been shown ^57^, and it is possible that elevated stress going along forced alcohol administration exacerbates the latent pro-inflammatory effects of alcohol exposure. Further research is required in order to better understand the relationship between alcohol exposure model, stress, induction of neuro-inflammation in the rodent PFC and the appearance of behavioral defects.

AAE has also been shown to disrupt neurotransmitter systems and reduce neurogenesis, by altering the expression of the brain-derived neurotrophic factor (BDNF) via changes in DNA methylation and/or acetylation ^14,38^. AAE-dependent alterations in epigenetic programming have also been reported and may be responsible for the delayed appearance of the long-lasting behavioral effects of AAE ^14,58^. Indeed, it is believed that alcohol affects epigenetic pathways, modifying chromatin remodeling, gene expression, dendritic spines morphology and synaptic plasticity, to ultimately affect neurocircuits function and behavior ^14,58^. Epigenetic and synaptic plasticity modulation by alcohol during adolescence could explain the progressive development of impaired behaviors observed in the present study, as well as the absence of AAE-induced behavioral defects in adolescent mice. Further research is required to unravel AAE-induced epigenetic reprogramming and its relationship with the delayed appearance of behavioral effects.

## Conclusions

This study shows that voluntary alcohol binge-drinking during adolescence leads to severe behavioral impairments in adult mice. Indeed, AAE severely increases anxiety-like behaviors, depressive-like behaviors and alcohol consumption in adulthood, while impairing recognition memory and behavioral flexibility. Although differences were noted between males and females, our data showed that AAE similarly affects their behaviors. Surprisingly, adolescent behaviors were not affected by alcohol binge-drinking, suggesting that AAE-dependent alteration of behaviors is a progressive and insidious process, whose consequences only emerge during adulthood. In this view, our findings are of great importance regarding the major public health issue that is adolescent binge-drinking, and could help refining the prevention strategies against harmful alcohol use in youth ^59,60^. Finally, we reported that in opposition with models of passive exposure to alcohol, voluntary binge-drinking in adolescent mice did not induce a major activation of neuro-inflammation in the PFC.

## Funding and disclosures

Laurent Nguyen is senior Research Associate from the F.R.S-F.N.R.S and Sophie Laguesse is Marie Sklodowska Curie Actions postdoctoral fellow. This work was supported by the European Union’s Horizon 2020 research and innovation program under the Marie Sklodowska-Curie grant agreement 839178 (S.L.), the Fondation Francqui (S.L.), the Brain and Behavior Research Foundation (S.L.), the Fonds Leon Fredericq (S.L. and L.N.), the F.R.S.-F.N.R.S. (Synet; EOS 0019118F-RG36), the Fondation Médicale Reine Elisabeth (L.N.), the Fondation Simone et Pierre Clerdent (L.N), and the foundation Marie-Marguerite Delacroix (L.V.H). The authors declare no competing financial interests.

## Acknowledgments

We thank Romain Le Bail for graphical design with BioRender. We are grateful to Prof. Dorit Ron for her constructive feedback on the manuscript as well as to all members of the Nguyen laboratory for their critical reading.

## Author contributions

Conceptualization (S.L., L.N. and L.V.H); Formal analysis (S.L. and V.D); Investigation (L.V.H., S.L., A.A., M.C.B., T.V.I., V.D.); Methodology (S.L., L.V.H., V.D., E.Q.); Funding acquisition (L.N. and S.L.); Supervision (L.N. and E.Q.); Writing – original draft (S.L. and L.V.H.); Writing-editing (L.N., V.D., E.Q.).

## Notes

### Competing Interest Statement

The authors have declared no competing interest.

## References

1. Giedd JN. The teen brain: insights from neuroimaging. J Adolesc Health. 2008;42(4):335–343.

2. Romer D, Reyna VF, Satterthwaite TD. Beyond stereotypes of adolescent risk taking: Placing the adolescent brain in developmental context. Dev Cogn Neurosci. 2017;27:19–34.

3. Casey BJ. Beyond simple models of self-control to circuit-based accounts of adolescent behavior. Annu Rev Psychol. 2015;66:295–319.

4. Volkow ND, Fowler JS, Wang GJ. The addicted human brain: insights from imaging studies. J Clin Invest. 2003;111(10):1444–1451.

5. Chwedorowicz R, Skarzynski H, Pucek W, Studzinski T. Neurophysiological maturation in adolescence - vulnerability and counteracting addiction to alcohol. Ann Agric Environ Med. 2017;24(1):19–25.

6. Crews FT, Vetreno RP, Broadwater MA, Robinson DL. Adolescent Alcohol Exposure Persistently Impacts Adult Neurobiology and Behavior. Pharmacol Rev. 2016;68(4):1074–1109.

7. Organization WH. Global status report on alcohol and health 2018. Geneva, 2018. 2018.

8. Rolland B, Naassila M. Binge Drinking: Current Diagnostic and Therapeutic Issues. CNS Drugs. 2017;31(3):181–186.

9. Alcoholism NIoAAa. NIAAA Council Approves Definition of Binge Drinking. NIAAA Newsl. 2004;3(3).

10. Squeglia LM, Tapert SF, Sullivan EV, et al. Brain development in heavy-drinking adolescents. Am J Psychiatry. 2015;172(6):531–542.

11. Spear LP. Effects of adolescent alcohol consumption on the brain and behaviour. Nat Rev Neurosci. 2018;19(4):197–214.

12. Cservenka A, Brumback T. The Burden of Binge and Heavy Drinking on the Brain: Effects on Adolescent and Young Adult Neural Structure and Function. Front Psychol. 2017;8:1111.

13. Lannoy S, Heeren A, Dormal V, Billieux J, Maurage P. Is there room for attentional impairments in binge drinking? A commentary on Carbia et al. (2018). Neurosci Biobehav Rev. 2019;98:58–60.

14. Kyzar EJ, Floreani C, Teppen TL, Pandey SC. Adolescent Alcohol Exposure: Burden of Epigenetic Reprogramming, Synaptic Remodeling, and Adult Psychopathology. Front Neurosci. 2016;10:222.

15. Lees B, Meredith LR, Kirkland AE, Bryant BE, Squeglia LM. Effect of alcohol use on the adolescent brain and behavior. Pharmacol Biochem Behav. 2020;192:172906.

16. Hiller-Sturmhofel S, Spear LP. Binge Drinking’s Effects on the Developing Brain-Animal Models. Alcohol Res. 2018;39(1):77–86.

17. Crews FT, Vetreno RP. Addiction, adolescence, and innate immune gene induction. Front Psychiatry. 2011;2:19.

18. Alfonso-Loeches S, Pascual M, Guerri C. Gender differences in alcohol-induced neurotoxicity and brain damage. Toxicology. 2013;311(1-2):27–34.

19. Pascual M, Blanco AM, Cauli O, Minarro J, Guerri C. Intermittent ethanol exposure induces inflammatory brain damage and causes long-term behavioural alterations in adolescent rats. Eur J Neurosci. 2007;25(2):541–550.

20. Guerri C, Pascual M. Impact of neuroimmune activation induced by alcohol or drug abuse on adolescent brain development. Int J Dev Neurosci. 2019;77:89–98.

21. Thiele TE, Crabbe JC, Boehm SL, 2nd. “Drinking in the Dark” (DID): a simple mouse model of binge-like alcohol intake. Curr Protoc Neurosci. 2014;68:9 49 41–12.

22. Laviola G, Macri S, Morley-Fletcher S, Adriani W. Risk-taking behavior in adolescent mice: psychobiological determinants and early epigenetic influence. Neurosci Biobehav Rev. 2003;27(1-2):19–31.

23. Laguesse S, Morisot N, Phamluong K, Ron D. Region specific activation of the AKT and mTORC1 pathway in response to excessive alcohol intake in rodents. Addict Biol. 2017;22(6):1856–1869.

24. Zapata A, Gonzales RA, Shippenberg TS. Repeated ethanol intoxication induces behavioral sensitization in the absence of a sensitized accumbens dopamine response in C57BL/6J and DBA/2J mice. Neuropsychopharmacology. 2006;31(2):396–405.

25. Fitzgerald PJ, Hale PJ, Ghimire A, Watson BO. The cholinesterase inhibitor donepezil has antidepressant-like properties in the mouse forced swim test. Transl Psychiatry. 2020;10(1):255.

26. Himanshu, Dharmila, Sarkar D, Nutan. A Review of Behavioral Tests to Evaluate Different Types of Anxiety and Anti-anxiety Effects. Clin Psychopharmacol Neurosci. 2020;18(3):341–351.

27. Kraeuter AK, Guest PC, Sarnyai Z. The Forced Swim Test for Depression-Like Behavior in Rodents. Methods Mol Biol. 2019;1916:75–80.

28. Leger M, Quiedeville A, Bouet V, et al. Object recognition test in mice. Nat Protoc. 2013;8(12):2531–2537.

29. Moy SS, Nadler JJ, Perez A, et al. Sociability and preference for social novelty in five inbred strains: an approach to assess autistic-like behavior in mice. Genes Brain Behav. 2004;3(5):287–302.

30. Riedel G, Robinson L, Crouch B. Spatial learning and flexibility in 129S2/SvHsd and C57BL/6J mouse strains using different variants of the Barnes maze. Behav Pharmacol. 2018;29(8):688–700.

31. Ron D, Barak S. Molecular mechanisms underlying alcohol-drinking behaviours. Nat Rev Neurosci. 2016;17(9):576–591.

32. Laguesse S, Morisot N, Shin JH, et al. Prosapip1-Dependent Synaptic Adaptations in the Nucleus Accumbens Drive Alcohol Intake, Seeking, and Reward. Neuron. 2017;96(1):145–159 e148.

33. Thiele TE, Navarro M. “Drinking in the dark” (DID) procedures: a model of binge-like ethanol drinking in non-dependent mice. Alcohol. 2014;48(3):235–241.

34. Kokare DM, Kyzar EJ, Zhang H, Sakharkar AJ, Pandey SC. Adolescent Alcohol Exposure-Induced Changes in Alpha-Melanocyte Stimulating Hormone and Neuropeptide Y Pathways via Histone Acetylation in the Brain During Adulthood. Int J Neuropsychopharmacol. 2017;20(9):758–768.

35. Pandey SC, Sakharkar AJ, Tang L, Zhang H. Potential role of adolescent alcohol exposure-induced amygdaloid histone modifications in anxiety and alcohol intake during adulthood. Neurobiol Dis. 2015;82:607–619.

36. Sakharkar AJ, Vetreno RP, Zhang H, Kokare DM, Crews FT, Pandey SC. A role for histone acetylation mechanisms in adolescent alcohol exposure-induced deficits in hippocampal brain-derived neurotrophic factor expression and neurogenesis markers in adulthood. Brain Struct Funct. 2016;221(9):4691–4703.

37. Antunes M, Biala G. The novel object recognition memory: neurobiology, test procedure, and its modifications. Cogn Process. 2012;13(2):93–110.

38. Crews FT, Robinson DL, Chandler LJ, et al. Mechanisms of Persistent Neurobiological Changes Following Adolescent Alcohol Exposure: NADIA Consortium Findings. Alcohol Clin Exp Res. 2019;43(9):1806–1822.

39. Zhang L, Zhang J, You Z. Switching of the Microglial Activation Phenotype Is a Possible Treatment for Depression Disorder. Front Cell Neurosci. 2018;12:306.

40. Vetreno RP, Crews FT. Adolescent binge drinking increases expression of the danger signal receptor agonist HMGB1 and Toll-like receptors in the adult prefrontal cortex. Neuroscience. 2012;226:475–488.

41. Montesinos J, Pascual M, Pla A, et al. TLR4 elimination prevents synaptic and myelin alterations and long-term cognitive dysfunctions in adolescent mice with intermittent ethanol treatment. Brain Behav Immun. 2015;45:233–244.

42. Montesinos J, Pascual M, Rodriguez-Arias M, Minarro J, Guerri C. Involvement of TLR4 in the long-term epigenetic changes, rewarding and anxiety effects induced by intermittent ethanol treatment in adolescence. Brain Behav Immun. 2016;53:159–171.

43. Coleman LG, Jr., Liu W, Oguz I, Styner M, Crews FT. Adolescent binge ethanol treatment alters adult brain regional volumes, cortical extracellular matrix protein and behavioral flexibility. Pharmacol Biochem Behav. 2014;116:142–151.

44. Lamont MG, McCallum P, Head N, Blundell J, Weber JT. Binge drinking in male adolescent rats and its relationship to persistent behavioral impairments and elevated proinflammatory/proapoptotic proteins in the cerebellum. Psychopharmacology (Berl). 2020;237(5):1305–1315.

45. Amodeo LR, Wills DN, Sanchez-Alavez M, Nguyen W, Conti B, Ehlers CL. Intermittent voluntary ethanol consumption combined with ethanol vapor exposure during adolescence increases drinking and alters other behaviors in adulthood in female and male rats. Alcohol. 2018;73:57–66.

46. Gass JT, Glen WB, Jr., McGonigal JT, et al. Adolescent alcohol exposure reduces behavioral flexibility, promotes disinhibition, and increases resistance to extinction of ethanol self-administration in adulthood. Neuropsychopharmacology. 2014;39(11):2570–2583.

47. Lee KM, Coehlo MA, Solton NR, Szumlinski KK. Negative Affect and Excessive Alcohol Intake Incubate during Protracted Withdrawal from Binge-Drinking in Adolescent, But Not Adult, Mice. Front Psychol. 2017;8:1128.

48. Pascual M, Lopez-Hidalgo R, Montagud-Romero S, Urena-Peralta JR, Rodriguez-Arias M, Guerri C. Role of mTOR-regulated autophagy in spine pruning defects and memory impairments induced by binge-like ethanol treatment in adolescent mice. Brain Pathol. 2021;31(1):174–188.

49. Al Shoyaib A, Archie SR, Karamyan VT. Intraperitoneal Route of Drug Administration: Should it Be Used in Experimental Animal Studies? Pharm Res. 2019;37(1):12.

50. Sabry FM, Ibrahim MK, Hamed MR, Ahmed HMS. Neurobehavioral effects of alcohol in overcrowded male adolescent rats. Neurosci Lett. 2020;731:135084.

51. Pearson BL, Defensor EB, Blanchard DC, Blanchard RJ. C57BL/6J mice fail to exhibit preference for social novelty in the three-chamber apparatus. Behav Brain Res. 2010;213(2):189–194.

52. Tambour S, Brown LL, Crabbe JC. Gender and age at drinking onset affect voluntary alcohol consumption but neither the alcohol deprivation effect nor the response to stress in mice. Alcohol Clin Exp Res. 2008;32(12):2100–2106.

53. Salguero A, Suarez A, Luque M, et al. Binge-Like, Naloxone-Sensitive, Voluntary Ethanol Intake at Adolescence Is Greater Than at Adulthood, but Does Not Exacerbate Subsequent Two-Bottle Choice Drinking. Front Behav Neurosci. 2020;14:50.

54. Spear LP. Consequences of adolescent use of alcohol and other drugs: Studies using rodent models. Neurosci Biobehav Rev. 2016;70:228–243.

55. Vetreno RP, Qin L, Crews FT. Increased receptor for advanced glycation end product expression in the human alcoholic prefrontal cortex is linked to adolescent drinking. Neurobiol Dis. 2013;59:52–62.

56. Holford NH. Clinical pharmacokinetics of ethanol. Clin Pharmacokinet. 1987;13(5):273–292.

57. Weera MM, Gilpin NW. Biobehavioral Interactions Between Stress and Alcohol. Alcohol Res. 2019;40(1).

58. Pandey SC, Kyzar EJ, Zhang H. Epigenetic basis of the dark side of alcohol addiction. Neuropharmacology. 2017;122:74–84.

59. Salmanzadeh H, Ahmadi-Soleimani SM, Pachenari N, et al. Adolescent drug exposure: A review of evidence for the development of persistent changes in brain function. Brain Res Bull. 2020;156:105–117.

60. Ryan SA, Kokotailo P, Committee On Substance USE, Prevention. Alcohol Use by Youth. Pediatrics. 2019;144(1).

