## Supplemental information for "Adolescent alcohol binge-drinking induces delayed appearance of behavioral defects in mice"

**Supplementary information**

**Supplemental Material and methods**

Blood Alcohol Concentration Measurement

Blood alcohol concentration (BAC) was measured as previously described ^1,2^. Immediately at the end of the last alcohol drinking session, mice were euthanized and trunk blood was collected in heparinized capillary tubes. Serum was extracted with 3.4% perchloric acid followed by centrifugation (5 minutes, 420g) and assayed for alcohol content using the nicotinamide adenine dinucleotide redox (NAD+/NADH) enzyme spectrophotometric method. BACs were determined by using a standard calibration curve.

Open field test

Locomotion and anxiety-like behavior were evaluated by using the Open Field (OF). Locomotor activity was used as a measure of overall activity, and the same experimental protocol was used two consecutive days to evaluate the locomotor activity and its habituation. On testing days, all mice were placed into the OF (40x40x40 cm, white Plexiglas walls and floor), illuminated by indirect white lightening by three 80 lux lamps. At the start of each trial, the mouse was gently placed in the center of the OF and allowed to walk freely. The distance travelled (cm) was recorded for 30 min by an activity tracking system, Videotracking® (France, Lyon). A second cohort of mice were used to assess thymotagsis in the OF. Time spent in the center of the OF was recorded on three consecutive days for 5 minutes. At the end of the trial, the mouse was immediately returned to its homecage.

Elevated Plus Maze Test

The EPM consists of two opposite open arms and two opposite opaque closed arms (29*20*6), which are connected by a square center zone. The maze was elevated 80cm from the floor and illuminated by three lamps (~80lux). Animals were placed in the center of the EPM and were allowed to freely explore the arms for 5 minutes. The time spent by the mouse in open/unprotected maze arms, closed/protected maze arms and in center was recorded. Time spent in the center of the maze was measured independently of closed arms time and was similar for all groups of mice (data not shown). The total number of entries in open and closed arms was used as a measure of the general activity ^3^.

Forced swimming Test

This experimental model was performed according to the procedures described by Porsolt *et al*. ^4^. Mice were forced to swim in a transparent glass cylinder (30 cm in height per 20 cm in diameter) filled with water to a 15-cm depth (25 °C ± 1◦C) and monitored for 6 min using a camera. The immobility time was determined during the last 4 min of the single experimental session. Mice were considered immobile when they ceased struggling and remained motionlessly floating in the water, making only movements necessary to keep their heads above the water surface. After the behavior assessment, the animals were placed on a heating bed until complete drying and then brought back to the cages.

Novel Object Recognition Test

NOR was performed in the OF using the protocol described by Leger *et al* ^5^, with a long habituation phase. Mice were placed in the empty OF once a day (10 minutes) for 3 consecutive days for habituation. Twenty-four hours after last habituation session, two identical objects were placed in the box (10 cm from the walls) and mice were allowed to familiarize with both objects for 10 minutes. Position of the objects in the OF was alternated between mice. The time for reaching 20 seconds of exploration was measured. Mice which did not reach the 20 seconds exploration criterion were excluded from the study (2/110 mice). Twenty-four hours after the familiarization session, the test session began. One object was replaced by another object (different in shape, color and texture), and mice were allowed to explore both objects for 10 minutes. Novel object was alternated between mice. Time to reach 20 seconds exploration was recorded and mice which did not reach the criterion were excluded from the study (6 /110 mice). The time spent exploring the familiar and the novel object was recorded. Valid exploration was characterized by the nose of the mouse directed toward an object at less than 2 cm of distance. Discrimination index was calculated as (Time exploring the novel object - Time exploring the familiar object) / Total exploration time. Familiar object habituation index was calculated as (Time exploring both objects during familiarization/2)-Time exploring familiar object during test session.

Three Chamber Test

The three chamber test was employed to characterize sociability and preference for social novelty, as described in Moy *et al* ^6^. Briefly, after 5 minutes of habituation in the apparatus, a stranger mouse was placed inside a wired cup. Position of the mouse and the empty wired cup were alterned between tested subjects. Mice were allowed to freely move between social and non-social chambers for 10 minutes. Sociability was evaluated by examining the amount of time spent by the mouse sniffing the social stimulus (i.e. the mouse) and the time spent sniffing the non-social stimulus (i.e. the empty wired cup). Then, a novel mouse was introduced in the empty wired cage, and mice were allowed to freely move between chambers for 10 minutes. Social novelty preference index (SNI) was calculated as (Time sniffing new mouse – Time sniffing familiar mouse) / (Time sniffing both mice). Position of the familiar and novel mouse was alternated between tested subjects.

Reversal Learning Test

The reversal learning test was performed by using the Barnes maze, as described in Riedel *et al* ^7^ and Pitts *et al* ^8^, with slight modifications. The maze consisted of an open circular area (120cm diameter) with 20 holes (5 cm diameter) regularly spaced, placed 2,5cm from the edge. The maze had no perimeter walls and was elevated 80cm from the floor. The arena was surrounded by four distal cues. Both direct and indirect illumination of the maze (~120 lx) and loud music (~90dB) provided an aversive stimulus to motivate escape. A black opaque target box containing a tissue paper was attached to the underside of the arena under one of the holes and was the only mean to escape from the bright arena. All sessions were recorded by an overhead camera.

Learning

Mice were first habituated to the escape box for 1 minute and to the Barnes maze platform for 5 minutes. Mice which did not enter the escape tunnel were directed toward it by the experimenter. One hour after the habituation, mice started the acquisition trainings. Training consisted of two acquisition trials daily (3 minutes limit per trial, intertrial interval of 1 hour). At the start of each trial, the mouse is placed in an opaque start chamber located in the center of the maze, which is lifted to allow the mouse exploring the maze. The position of the escape tunnel remained the same during the acquisition training. For each trial, a blinded experimenter recorded the delay before entering the escape tunnel, as well as the number of primary errors (number of incorrect holes checked by the mouse). Seventy-two hours after the last learning session, mice underwent a 80-second probe trial, in which the escape tunnel was removed from the apparatus. During the probe trial, time spent in the correct sector (in the quadrant containing the correct hole) was recorded.

Reversal learning

Twenty-four hours after the learning probe trial, mice underwent 10 sessions of reversal learning (2 sessions per day, intertrial interval of 1 hour), in which the escape tunnel location was relocated to the hole that was exactly 180° from the initial location. Time to escape and number of primary errors were analyzed. Seventy-two hours after the last reversal learning session, mice underwent a probe test, in which time spent in the correct sector and in the sector of the previous location of the escape tunnel were recorded.

Two-bottle choice drinking paradigm

Intermittent Access to 20% alcohol

The intermittent access to 20% alcohol two-bottle choice drinking procedure (IA20%-2BC) was conducted as previously described ^9,10^ (Fig 5A). Mice were single-housed and weighted every other day. Mice were given 24 hours of concurrent access to one bottle of 20% alcohol (v/v) in tap water, and one bottle of water. Drinking sessions started at 12:00 on Monday, Wednesday and Friday, with 24- or 48 hours (weekend) of alcohol-deprivation periods in which mice consumed only water. The placement (left or right) of water and alcohol bottle was alternated between each session in order to control for side preference. Water and alcohol bottles were weighted at the beginning of each alcohol session and at the end of each alcohol sessions.

Intermittent Access to 1% sucrose

Mice underwent an IA-2BC paradigm as described above but had access to one bottle containing 1% sucrose solution in water, and another bottle containing water.

**Immunohistochemistry**

Immunohistochemistry was conducted as previously described ^11^. Fifty µm brain sections were incubated in the primary antibody overnight at 4C (Goat anti-Iba1 1/500, abcam (Cambridge, UK) #ab5076). Donkey anti-goat AlexaFluor 564 was used as secondary antibody for 4 hours at room temperature, with DAPI. Images were acquired on a Nikon A1 confocal microscope and NIS-Element Imaging software at 20x magnification. Prelimbic and infralimbic medial sections (636,4µm x 636,4µm) were imaged in z-stack (1µm, 7-11 images per stack) and analyzed by blinded observer. Microglial cell activation state was defined according to their shape: microglia presenting a small nucleus and numerous and ramified extensions were considered as resting, whereas amoeboid-shaped microglial cells with a large nucleus and smaller extensions were considered as activated.

**ELISA**

Prefrontal cortices were homogenized in tissue lysis buffer (20 mM Tris, 150 mM NaCl, 1% Nonidet P-40, 0.5% sodium deoxycholate, 0.1% SDS, 50mM Tris, protease inhibitors cocktail; pH8) and centrifuged for 10 min at 10 000g at 4C. Supernatants were collected and protein concentrations were determined by using the BCA assay kit (Thermofischer Scientifics, Waltham, MA, USA). IL-1β levels were determined by using the IL-1β ELISA kit (ThermoFisher Scientific, #88-7013) following the manufacturer’s protocols.

**Supplemental Tables**

**Table S1: Individual alcohol-drinking data of mice used for behavioral experiments**

| Behavioral test | Sex | Age at test | Number | Mean alcohol consumption (g/kg/4h) |
| --- | --- | --- | --- | --- |
| Locomotion | Male | P43 | 8636; 8637; 8638; 8639; 8640; 8641; 8642; 8643; 8644; 8645; 8646; 8647 | 7.35; 8.68; 4.79; 7.69; 6.18; 9.61; 6.27; 8.04; 7.76; 7.94; 6.64; 6.09 Mean 7.25±1.3 |
|  | Female | P43 | 8626; 8627; 8628; 8629; 8630; 8631; 8632; 8633; 8634; 8635 | 8.21; 6.08; 8.01; 7.41; 5.48; 9.15; 8.04; 7.40; 8.06; 8.83 Mean 7.67±1.1 |
|  | Male | P80-100 | 6363; 6364; 6365; 6366; sb; 6368; 6369 | 7.15; 7.15; 5.85; 5.35; 7.37; 6.22; 5.79 Mean 6.41±0.8 |
|  | Female | P80-100 | 6352; 6353; 6354; 6355; 6356; 6357; 6358; 6361; 6362 | 7.51; 6.62; 8.46; 6.49; 8.01; 8.72; 6.83; 6.90; 7.03 Mean 7.4±0.82 |
| Open Field | Male | P43 | 4181; 4176; 4178; 4179; 4173; 4183; 4177; 1932; 1933; 1934; 1935; 1936; 8672; 8671 | 6.36; 6.17; 6.24; 5.15; 5.83; 5.26; 6.40; 5.63; 4.86; 8.63; 6.69; 6.70; 7.41; 8.19 Mean 6.39±1.1 |
|  | Female | P43 | 4169; 4164; 4167; 4163; 4170; 1937; 1939; 1940; 1941; 1942; 1944; 1957; 8677; 8675 | 7.32; 8.77; 5.55; 5.21; 7.34; 8.21; 4.86; 4.59; 7.43; 7.41; 7.56; 9.21; 8.22; 6.18 Mean 6.99±1.5 |
|  | Male | P80-100 | 3517; 3521; 3528; 3518; 3515; 3522; 3520; 1917; 1918; 1921; 1502; 1504 | 5.88; 5.07; 6.86; 5.69; 5.95; 6.58; 6.78; 6.37; 6.07; 7.37; 5.32; 6.1 Mean 6.17±0.62 |
|  | Female | P80-100 | 3536; ss bc; 3533; 3543; 3539; 3537; 3534; 1913; 1911; 1909; 1506; 1507 | 7.24; 6.87; 5.47; 6.65; 6.53; 6.65; 6.75; 7.08; 7.09; 6.53; 5.77; 6.78 Mean 6.62±0.52 |
| Elevated Plus Maze | Male | P43 | 7537; 7540; 7542; 7543; 7545; 1968; 1971; 1970; 1973; 1974; 1972; 1975 | 7.32; 5.08; 7.17; 4.48; 4.83; 8.27; 8.77; 7.89; 6.62; 7.07; 7.28; 6.62 Mean 6.78±1.36 |
|  | Female | P43 | 7550; 7561; 7551; 7549; 7563; 7564; 1955; 1953; 1954; 1956; 1957; 1959 | 6.35; 7.20; 5.22; 6.04; 8.23; 6.12; 8.15; 6.89; 8.85; 11.05; 10.96; 6.10 Mean 7.60±1.9 |
|  | Male | P80-100 | 3517; 3521; 3528; 3518; 3515; 3522; 3520; 1917; 1918; 1921; 1502; 1504; 6100 | 5.88; 5.07; 6.86; 5.69; 5.95; 6.58; 6.78; 6.37; 6.07; 7.37; 5.32; 6.1; 6.13 Mean 6.16±0.63 |
|  | Female | P80-100 | 3536; ss bc; 3533; 3543; 3539; 3534; 1913; 1911; 1909; 1506; 1507; 1508; 1509 | 7.24; 6.87; 5.47; 6.65; 6.53; 6.75; 7.08; 7.09; 6.53; 5.77; 6.78; 6.16; 6.53 Mean 6.57±0.51 |
| Forced Swimming Test | Male | P43 | 4103; 4104; 4101; 3000; 4105; 4106; 2999; 1778; 1781; 1762; 1763; 1784 | 6.34; 5.29; 6.59; 5.66; 4.31; 6.87; 4.89; 5.8; 7; 6.2; 5.3; 5.5 Mean 5.81±0.82 |
|  | Female | P43 | 4112; 4110; 4111; 4109; 4108; 1773; 1772; 1768; 1766; 1774; 1760 | 5.61; 8.71; 7.77; 4.56; 8.77; 7.3; 6; 5.6; 6.1; 7.4; 6.3 Mean 6.78±1.35 |
|  | Male | P80-100 | 6683; 6688; 6685; 6696; 6698; 7545; 7540; 7537; 1501; 1503; 6100; 1525; 1523; 1526; 1522; 1527 | 4.75; 6.90; 4.54; 5.41; 5.26; 4.83; 5.08; 7.32; 7.65; 5.93; 6.13; 6.38; 5.04; 5.45; 7.68; 7.39 Mean 5.99±1.1 |
|  | Female | P80-100 | 6690; 6692; 6687; 6686; 6693; 7550; 7563; 7561; 7549; 1511; 1510; 1508; 1536; 1534; 1535; 1537; 1533 | 8.11; 6.96; 6.82; 8.25; 7.60; 6.35; 8.23; 7.20; 6.04; 8.75; 7.43; 6.16; 5.21; 7.60; 8.93; 8.77; 7.34 Mean 7.40±1.1 |
| Novel Object Recognition Test | Male | P43 | 1932; 1935; 1936; 8671; 8672; 8673; 8668; 8669; 8670 | 5.63; 6.69; 6.70; 8.19; 7.41; 6.46; 7.87; 7.60; 5.60 Mean 6.91±0.93 |
|  | Female | P43 | 1937; 1939; 1940; 1941; 1942; 1944; 1957; 1943; 1938; 8674; 8676; 8678; 8679; 8680; 8681 | 8.21; 4.86; 4.59; 7.43; 7.41; 7.56; 9.21; 7.17; 5.43; 9.43; 10.00; 5.93; 9.18; 7.60; 8.50 Mean 7.5±1.7 |
|  | Male | P80-100 | 9772; 9773; 9774; 9771; 1917; 1918; 1504; 1502; 1503; 6100; 1501; 1523; 1524; 1525; 1526; 1527 | 7.09; 5.87; 8.62; 5.79; 6.37; 6.07; 6.10; 5.32; 5.93; 6.13; 7.65; 5.04; 5.44; 6.38; 5.45; 7.39 Mean 6.29±0.95 |
|  | Female | P80-100 | 9775; 9776; 9780; 9778; 9777; 9779; 1909; 1911; 1913; 1506; 1509; 1510; 1511; 1534; 1533; 1535; 1537 | 5.96; 7.76; 6.32; 4.78; 7.71; 6.42; 6.53; 7.09; 7.08; 5.77; 6.53; 7.43; 8.75; 7.60; 7.34; 8.93; 8.77 Mean 7.1±1.1 |
| Sociability/Social Novelty | Male | P43 | 2; 3; 4; 5; 6; 7; 8; 9; 11; 12; 13 | 5.68; 9.52; 8.87; 9.13; 4.72; 8.96; 8.06; 8.35; 7.33; 5.49; 8.12 Mean 7.6±1.6 |
|  | Female | P43 | 8682; 8683; 8684; 8685; 8686; 8687; 8688; 8689; 8690; 8691 | 6.3; 5.4; 7.1; 6.2; 9.1; 10.7; 5.2; 10.6; 10.3; 8.6 Mean 7.9±1.9 |
|  | Male | P80-100 | 1570; 1572; 1573; 1574; 1578; 1579; 1580; 1581; 1582; 1593; 1594; 1595; 1596; 1597; 1598 | 6.98; 5.85; 6.28; 6.85; 6.00; 7.71; 4.72; 5.99; 6.45; 6.48; 5.26; 5.36; 5.41; 4.93; 5.48 Mean 5.98±0.82 |
|  | Female | P80-100 | 1583; 1584; 1585; 1586; 1587; 1588; 1589; 1590; 1591; 1592; 1599; 1600; 1701; 1702; 1705 | 7.00; 8.67; 8.22; 5.16; 9.69; 10.27; 7.48; 7.42; 7.42; 7.73; 8.57; 6.88; 7.86; 5.96; 8.10 Mean 7.76±1.3 |
| Barnes Maze Test | Male | P43 | 2492; 2493; 2494; 2495; 2496; 979; 980; 981; 982; 983; 985; 986 | 8.83; 5.87; 7.88; 5.51; 7.60; 7.21; 4.63; 6.03; 8.45; 5.95; 6.81; 8.95 Mean 6.98±1.4 |
|  | Female | P43 | 2481; 2482; 2483; 2484; 2485; 2486; 992; 993; 994; 995; 996; 997 | 7.10; 8.43; 8.13; 6.77; 7.56; 7.52; 6.82; 6.86; 5.96; 7.00; 7.77; 9.75 Mean 7.47±0.98 |
|  | Male | P80-100 | 1582; 1580; 1581; 1572; 1570; 1579; 1575; 1571; 1574; 1595; 1573; 1596; 1578 | 6.45; 4.72; 5.99; 5.85; 6.98; 7.71; 6.43; 6.83; 6.85; 5.36; 6.28; 5.41; 6 Mean 6.22±0.79 |
|  | Female | P80-100 | 1701; 1600; 1702; 1588; 1585; 1592; 1586; 1591; 1584; 1590; 1583; 1589; 1587 | 7.86; 6.88; 5.96; 10.27; 8.22; 7.73; 5.16; 7.42; 8.67; 7.42; 7; 7.48; 9.69 Mean 7.67±1.4 |
| IA-20% 2BC Test | Male | P43 | 5363; 5339; 5338; 5367; 5341; 5340; 5365; 5366; 6612; 6611; 6610; 6609 | 5.46; 5.39; 4.71; 6.43; 7.59; 3.84; 6.73; 4.35; 7.00; 7.06; 6.00; 9.13 Mean 6.14±1.5 |
|  | Female | P43 | 5333; 5332; 5336; 5369; 5368; 5331; 5335; 8608; 8606; 8604; 8607; 8605 | 6.24; 6.69; 6.04; 6.87; 7.88; 5.78; 8.28; 8.36; 8.58; 5.05; 8.29; 8.85 Mean 7.24±1.3 |
|  | Male | P80-100 | 3053; 3054; 3055; 3065; 3047; 3048; 3049; 3050; 3051; 3052; 7542; 7543 | 6.52; 8.65; 5.93; 4.76; 5.30; 4.68; 5.60; 6.09; 5.41; 4.22; 7.16; 4.47 Mean 5.73±1.3 |
|  | Female | P80-100 | 3057  3058  3059  3060  3067  3069  3070  3071  3072  3073  7551  7565 | 7.28  7.50  7.66  7.98  4.70  6.03  4.89  5.84  6.61  5.34  5.22  5.96  Mean 6.25±1.1 |

**Table S2: Individual alcohol-drinking data of mice used for biochemical experiments**

| Experiment | Sex | Age | Number | Mean alcohol consumption (g/kg/4h) |
| --- | --- | --- | --- | --- |
| Immunofluorescence | Male | P43 | 987  988  989  991 | 5.6  5.1  8.8  6.9  Mean 6.6±1.25 |
|  | Female | P43 | 303  304  305  307 | 9.3  5.3  4.9  7.5  Mean 6.75 ±1.65 |
|  | Male | P80 | 1969  1970  1971  1975 | 8.5  7.9  8.8  6.6  Mean 7.95±0.7 |
|  | Female | P80 | 1954  1955  1958  1959 | 8.8  8.2  8.5  6.1  Mean 7.9 ±0.9 |
| Western blot | Male | P43 | 9975  9972  9963  9961 | 8.6  10.2  7.8  8.1  Mean 8.6±0.7 |
|  | Female | P43 | 9946  9949  9954  9953 | 8.6  7.8  10.4  7.0  Mean 8.44±1 |
|  | Male | P80 | 9964  9965  9974  9977 | 7.9  6.0  5.5  6.6  Mean 6.5±0.7 |
|  | Female | P80 | 9957  9958  9956  9959 | 9.9  6.1  8.3  7.3  Mean 7.9±1.2 |
| ELISA | Male | P43 | 6428  6430  6431  6432 | 5.6  5.4  7.3  8.1  Mean 6.6±1.1 |
|  | Female | P43 | 6439  6441  6442  6443 | 8.7  9.5  11.5  8.0  Mean 9.4±1.1 |
|  | Male | P43 | 6429  6433  9970  9971 | 5.2  7.9  6.7  6.3  Mean 6.5±0.7 |
|  | Female | P80 | 6444  6445  6446  5595 | 5.7  5.3  5.5  7.5  Mean 6.02±0.8 |

**Supplemental Figure legends**

**Figure S1**: **Adolescent alcohol exposure does not alter locomotion activity or habituation**

**(A)** Short-term; Mean locomotion activity in the Open Field (OF). Adolescent mice were allowed to explore the OF for 30 minutes and the test was repeated 24 hours later. Locomotor activity (cm) was measured every 5 minutes. Linear mixed-effects model showed a significant main effect of interval (χ^2^_(5)_=507.7, p<0.001; η^2^=0.38) and day (χ^2^_(1)_=41.7, p<0.001, η^2^=0.13) but no effect of treatment (χ^2^_(1)_=1.1, p=0.29) or sex (χ^2^_(1)_=0, p=0.96); n= 9-12 per group. **(B)** Short-term; Total number of arm entries in the EPM. Two-way ANOVA showed no main effect of treatment (F_(1,46)_=2.18, p=0.15) or sex (F_(1,46)_=0.07, p=0.8); n=12-14 per group. **(C)** Long-term; Mean locomotion activity in the Open Field (OF). Linear mixed-effects model showed a significant main effect of interval (χ^2^_(5)_=610.6, p<0.001, η^2^=0.41), day (χ^2^_(1)_=66.18, p<0.001, η^2^=0.19) and sex (χ^2^_(1)_=4.81, p<0.05, η^2^=0.07) but no effect of treatment (χ^2^_(1)_=0.01, p=0.93); n=7-10 per group. **(D)** Long-term; Total number of arm entries in the EPM. Two-way ANOVA showed no main effect of treatment (F_(1,50)_=0.73, p=0.4) or sex (F_(1,50)_=1.23, p=0.27); n=13-14 per group.

**Figure S2: AAE does not impair sociability**

(A) Schematic representation of the three-chamber sociability test. The experimental apparatus consists in three chambers connected via sliding doors. For sociability assessment, an empty wire cage is placed in the non-social chamber and a cage containing a mouse is placed in the social chamber (left scheme). For social novelty assessment, a new mouse is introduced in the empty wire cage (right scheme). **(B)** Short-term; Sociability: percentage of time sniffing the mouse and the empty wire cage. Linear mixed-effects model showed a significant main effect of stimulus (mouse vs empty cage χ^2^_(1)_=86.37, p<0.001, η^2^=0.67) but no effect of sex (χ^2^_(1)_=2.76, p=0.1) or treatment (χ^2^_(1)_=0.12, p=0.72) and no interaction (sex x treatment (χ^2^_(1)_=0.9, p=0.34), sex x stimulus (χ^2^_(1)_=0.0, p=0.98), treatment x stimulus (χ^2^_(1)_=2.03, p=0.15); n= 10-12 per group. **(C)** Short-term; Social Novelty: social novelty preference index (SNI) was calculated as (Time sniffing new mouse – Time sniffing familiar mouse) / (Time sniffing both mice). Two-way ANOVA showed a significant main effect of sex (F_(1,39)_=4.1, p<0.05) but no main effect of treatment (F_(1,39)_=0.03, p=0.85) and no interaction (F_(1,39)_=0.07, p=0.79); n= 15-16 per group. **(D)** Long-term; Sociability: percentage of time sniffing the mouse and the empty wired cage. Linear mixed-effects model showed a significant main effect of stimulus (mouse vs empty cage χ^2^_(1)_=70.27, p<0.001, η^2^=0.54) but no effect of sex (χ^2^_(1)_=3.58, p=0.06) or treatment (χ^2^_(1)_=1.48, p=0.22) and no interaction (sex x treatment (χ^2^_(1)_=0.01, p=0.91), sex x stimulus (χ^2^_(1)_=0.17, p=0.68), treatment x stimulus (χ^2^_(1)_=0.02, p=0.89); n=10-12 per group. **(E)** Long-term; Social Novelty: social novelty preference index. Two-way ANOVA showed no main effect of sex (F_(1.57)_=1.27, p=0.26) or treatment (F_(1,57)_=0.41, p=0.52) and no interaction (F_(1,57)_=0.19, p=0.66); n=15-16 per group.

**Figure S3: AAE does not induce spatial learning defects.**

**(A)** Short-term; Learning: mean escape time per day (mean of two sessions) across 5 days. Linear mixed-effects model showed a significant main effect of day (χ^2^_(4)_=753.1, p<0.001, η^2^=0.76) but no effect of sex (χ^2^_(1)_=0.24, p=0.62) or treatment (χ^2^_(1)_=0.03, p=0.85) and no interaction (day x sex χ^2^_(4)_=2.20, p=0.7; day x treatment χ^2^_(4)_=0.43, p=0.98; sex x treatment χ^2^_(1)_=0.03, p=0.86). **(B)** Short-term; Learning: primary errors per day (mean of two sessions). Linear mixed-effects model showed a significant main effect of day (χ^2^_(4)_=351.43, p<0.001, η^2^=0.59), but no effect of sex (χ^2^_(1)_=0.84, p=0.36) or treatment (χ^2^_(1)_=2.92, p=0.09) and no interaction (day x sex χ^2^_(4)_=4.71, p=0.32; day x treatment χ^2^_(4)_=1.09, p=0.9; sex x treatment χ^2^_(1)_=0.02, p=0.88). **(C)** Short-term; Percentage of time spent in the correct sector during probe test (72 hours after the last learning session). Two-way ANOVA showed no main effect of treatment (F_(1,44)_=0.01, p=0.91) or sex (F_(1,44)_=1.15, p=0.29) and no interaction (F_(1,44)_=0.62, p=0.44); n=12 per group. **(D)** Long-term; Learning: mean escape time per day. Linear mixed-effects model showed a significant main effect of day (χ^2^_(4)_=683.02, p<0.001, η^2^=0.71) but no effect of sex (χ^2^_(1)_=1.43, p=0.23) or treatment (χ^2^_(1)_=0.15, p=0.7) and no interaction (day x sex χ^2^_(4)_=6, p=0.2; day x treatment χ^2^_(4)_=0.91, p=0.92; sex x treatment χ^2^_(1)_=0.02, p=0.87). **(E)** Long-term; Learning: primary errors number. Linear mixed-effects model showed a significant main effect of day (χ^2^_(4)_=152.26, p<0.001, η^2^=0.35), but no effect of sex (χ^2^_(1)_=0.13, p=0.72) or treatment (χ^2^_(1)_=0.21, p=0.65) and no interaction (day x sex χ^2^_(4)_=2.81, p=0.59; day x treatment χ^2^_(4)_=1.56, p=0.82; sex x treatment χ^2^_(1)_=0.02, p=0.9). **(F)** Long-term; Percentage of time spent in the correct sector during probe test (72 hours after the last learning session). Two-way ANOVA showed no main effect of treatment (F_(1,48)_=0.60, p=0.44) or sex (F_(1,48)_=2.5, p=0.12) and no interaction (F_(1,48)_=0.34, p=0.56); n= 13 per group.

**Figure S4: Adolescent alcohol exposure does not increase sucrose consumption.**

**(A)** Short-term; Sucrose intake (ml/kg/24h). Linear mixed-effects model showed a significant main effect of session (χ^2^_(4)_=149.38, p<0.001, η^2^=0.35) and sex (χ^2^_(1)_=4.78, p<0.05, η^2^=0.09) but no effect of treatment (χ^2^_(1)_=0.24, p=0.62) and no interaction (session x treatment χ^2^_(4)_=6.88, p=0.14, sex x treatment χ^2^_(1)_=0.53, p=0.47, session x sex χ^2^_(4)_=1.74, p=0.78). **(B)** Short-term; Sucrose preference. Linear mixed-effects model showed a significant main effect of session (χ^2^_(4)_=141.31, p<0.001, η^2^=0.34), but no effect of sex (χ^2^_(1)_=1.39, p=0.24) or treatment (χ^2^_(1)_=0.26, p=0.61) and no interaction (session x treatment χ^2^_(4)_=6.65, p=0.16, sex x treatment χ^2^_(1)_=0.16, p=0.69, session x sex χ^2^_(4)_=3.37, p=0.5). **(C)** Short-term; Total fluid intake (ml/kg/24h). Linear mixed-effects model showed a significant main effect of session (χ^2^_(4)_=57.78, p<0.001, η^2^=0.18) and sex (χ^2^_(1)_=6.02, p<0.01, η^2^=0.11) but no effect of treatment (χ^2^_(1)_=0.38 p=0.54) and no interaction (session x treatment χ^2^_(4)_=4.60, p=0.33, sex x treatment χ^2^_(1)_=1.29, p=0.26, s x sex χ^2^_(4)_=0.53, p=0.97); n=12 per group. **(D)** Long-term; Sucrose intake (ml/kg/24h). Linear mixed-effects model showed a significant main effect of session (χ^2^_(4)_=33.9, p<0.001, η^2^=0.12) and sex (χ^2^_(1)_=45.30, p<0.001, η^2^=0.2) but no effect of treatment (χ^2^_(1)_=0.01 p=0.93). This model also showed an interaction session x sex (χ^2^_(4)_=12.14, p<0.05, η^2^=0.04) but not sex x treatment (χ^2^_(1)_=0.07, p=0.79) or session x treatment (χ^2^_(4)_=6.26, p=0.18). **(E)** Long-term; Sucrose preference. Linear mixed-effects model showed a significant main effect of session (χ^2^_(4)_=39.4, p<0.001, η^2^=0.14) and sex (χ^2^_(1)_=5.58, p<0.05, η^2^=0.03) but no effect of treatment (χ^2^_(1)_=1.17, p=0.28) and no interaction (session x treatment χ^2^_(4)_=3, p=0.56, sex x treatment χ^2^_(1)_=0.86, p=0.35, session x sex χ^2^_(4)_=3.04, p=0.55). **(F)** Long-term; Total fluid intake (ml/kg/24h). Linear mixed-effects model showed a significant main effect of session (χ^2^_(4)_=12.34, p<0.01, η^2^=0.04) and sex (χ^2^_(1)_=65.14, p<0.001, η^2^=0.25) but no effect of treatment (χ^2^_(1)_=0.39 p=0.53). This model also showed an interaction session x Sex (χ^2^_(4)_=19.57, p<0.001, η^2^=0.07) but not sex x treatment (χ^2^_(1)_=0.59, p=0.44) or session x treatment (χ^2^_(4)_=4.99, p=0.29); n=12 per group.

**Figure S5: Voluntary alcohol binge-drinking during adolescence does not lead to long-lasting neuroinflammation in the prefrontal cortex**

**(A,B)** representative images of resting (**A**) or activated (**B**) microglial cells. Microglial cells with small nucleus, numerous ramified extensions and cell body to cell size ratio between 8 and 12 were considered as resting ^12^ (**A**), whereas amiboid-shaped microglial cells with a large nucleus, smaller extensions, cell body to cell size ratio between 12 and 20 were considered as activated (**B**). **(C)** Short-term; Total number of microglial cells per prelimbic PFC section (mean ± S.E.M.). Two-way ANOVA showed no main effect of treatment (F_(1,12)_=1.87, p=0.2), sex (F_(1,12)_=0.002, p=0.97) and no interaction (F_(1,12)_=1.27, p=0.28); **(D-G)** Long-term, Immunofluorescence analysis of microglial cells expressing Iba1 (red) in the prelimbic prefrontal cortex of males water (**D**), males AAE (**E**), females water (**F**) and females AAE (**G**); Scale bar 10µm. **(H)** Long-term; Number of activated Iba1+ microglial cells per prelimbic PFC section (mean +- S.E.M.). Two-way ANOVA showed no main effect of treatment (F_(1,12)_=0, p=0.99), sex (F_(1,12)_=1.8, p=0.19) and no interaction (F_(1,12)_=0, p=0.99). **(I)** Long-term; Total number of microglial cells per prelimbic PFC section (mean + S.E.M.). Two-way ANOVA showed no main effect of treatment (F_(1,12)_=0.04, p=0.83), sex (F_(1,12)_=1.4, p=0.27) and no interaction (F_(1,12)_=0.17, p=0.69); n=4 per group.

1.	Neasta J, Ben Hamida S, Yowell Q, Carnicella S, Ron D. Role for mammalian target of rapamycin complex 1 signaling in neuroadaptations underlying alcohol-related disorders. *Proc Natl Acad Sci U S A.* 2010;107(46):20093-20098.

2.	Zapata A, Gonzales RA, Shippenberg TS. Repeated ethanol intoxication induces behavioral sensitization in the absence of a sensitized accumbens dopamine response in C57BL/6J and DBA/2J mice. *Neuropsychopharmacology.* 2006;31(2):396-405.

3.	Himanshu, Dharmila, Sarkar D, Nutan. A Review of Behavioral Tests to Evaluate Different Types of Anxiety and Anti-anxiety Effects. *Clin Psychopharmacol Neurosci.* 2020;18(3):341-351.

4.	Porsolt RD, Le Pichon M, Jalfre M. Depression: a new animal model sensitive to antidepressant treatments. *Nature.* 1977;266(5604):730-732.

5.	Leger M, Quiedeville A, Bouet V, et al. Object recognition test in mice. *Nat Protoc.* 2013;8(12):2531-2537.

6.	Moy SS, Nadler JJ, Perez A, et al. Sociability and preference for social novelty in five inbred strains: an approach to assess autistic-like behavior in mice. *Genes Brain Behav.* 2004;3(5):287-302.

7.	Riedel G, Robinson L, Crouch B. Spatial learning and flexibility in 129S2/SvHsd and C57BL/6J mouse strains using different variants of the Barnes maze. *Behav Pharmacol.* 2018;29(8):688-700.

8.	Pitts MW. Barnes Maze Procedure for Spatial Learning and Memory in Mice. *Bio Protoc.* 2018;8(5).

9.	Ron D, Barak S. Molecular mechanisms underlying alcohol-drinking behaviours. *Nat Rev Neurosci.* 2016;17(9):576-591.

10.	Laguesse S, Morisot N, Phamluong K, Ron D. Region specific activation of the AKT and mTORC1 pathway in response to excessive alcohol intake in rodents. *Addict Biol.* 2017;22(6):1856-1869.

11.	Laguesse S, Morisot N, Shin JH, et al. Prosapip1-Dependent Synaptic Adaptations in the Nucleus Accumbens Drive Alcohol Intake, Seeking, and Reward. *Neuron.* 2017;96(1):145-159 e148.

12.	Hovens IB, van Leeuwen BL, Nyakas C, Heineman E, van der Zee EA, Schoemaker RG. Postoperative cognitive dysfunction and microglial activation in associated brain regions in old rats. *Neurobiol Learn Mem.* 2015;118:74-79.
