## Supplementary figures and images for "Adolescent alcohol binge-drinking induces delayed appearance of behavioral defects in mice"

### Supplemental figures

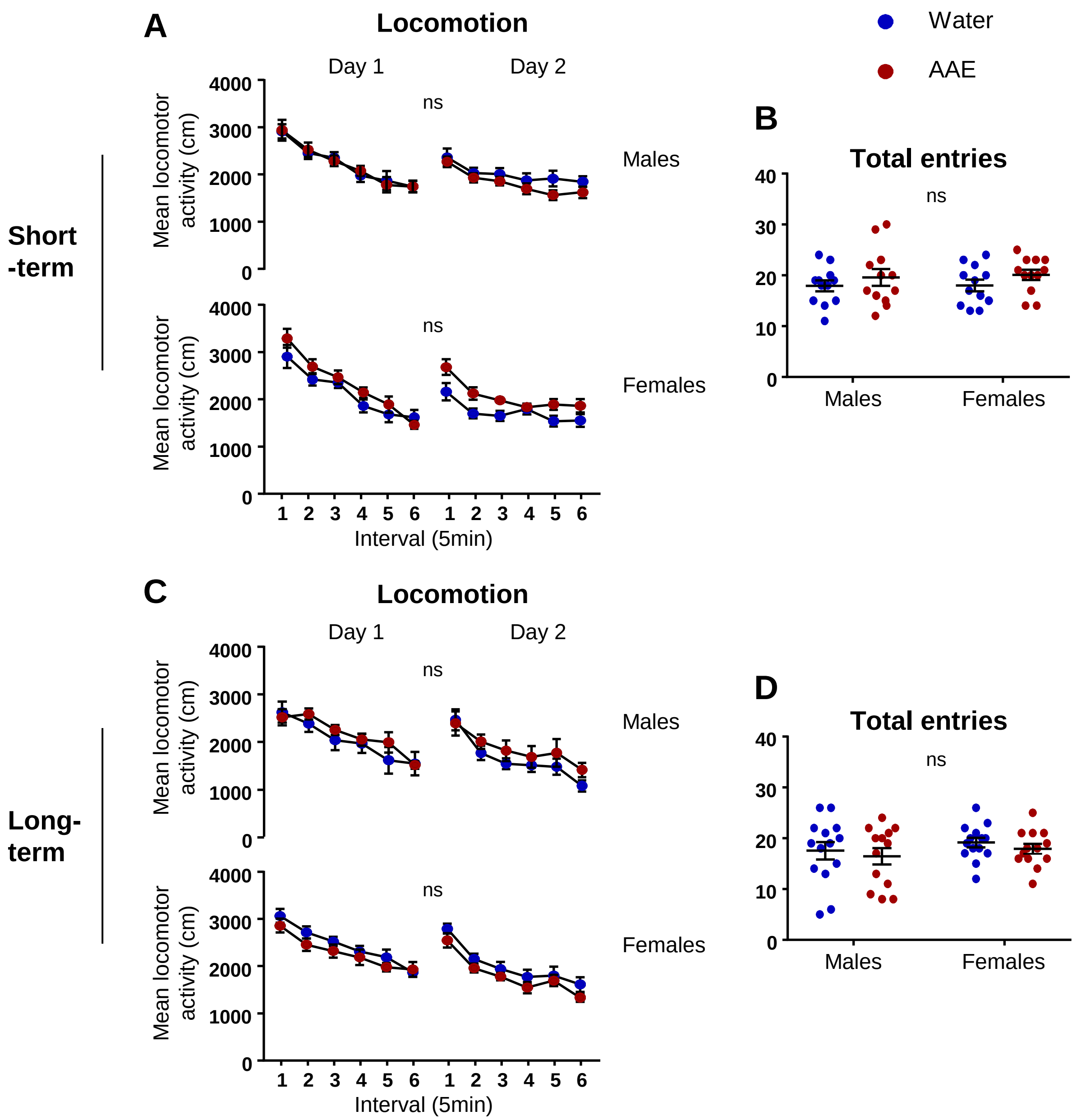

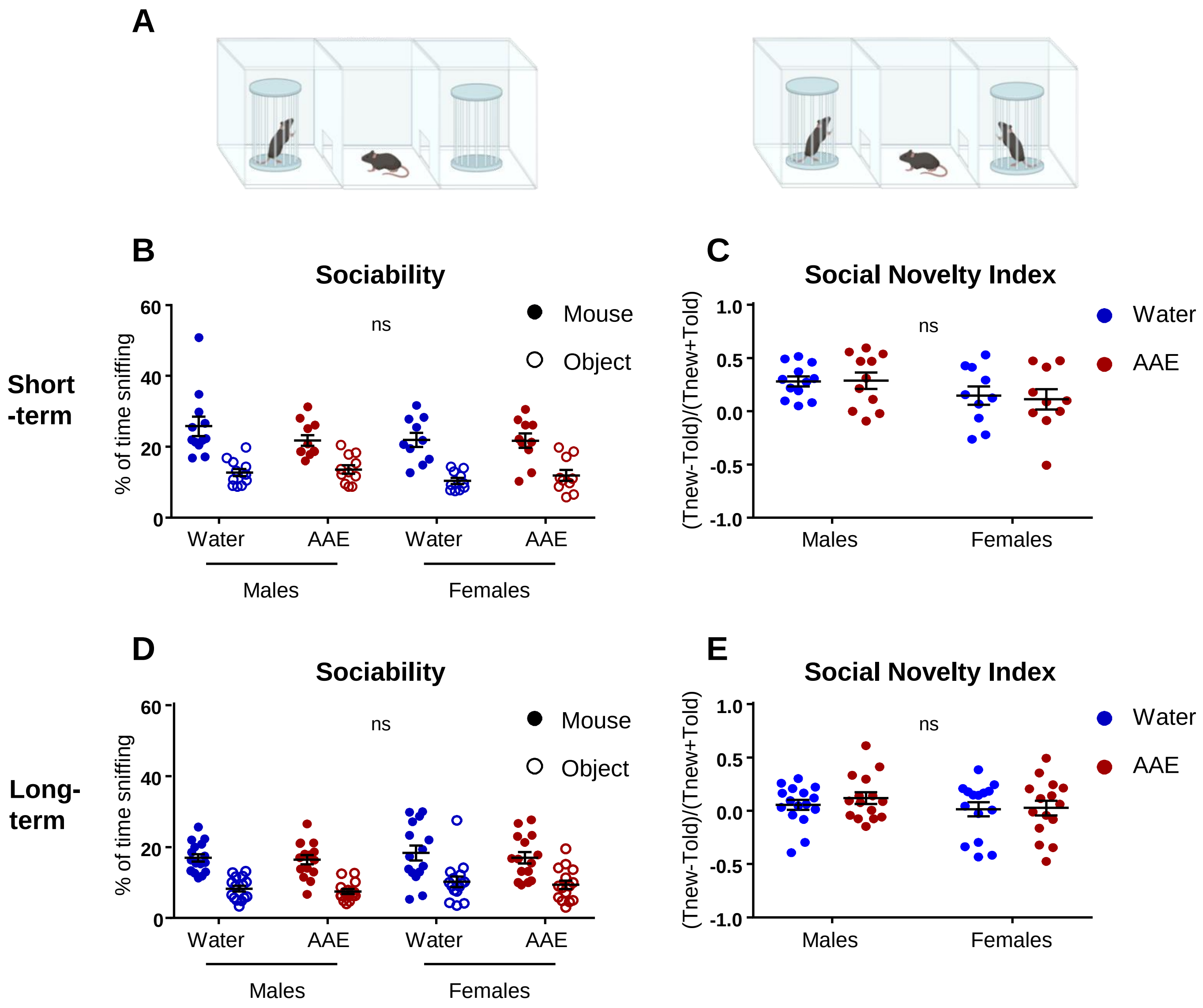

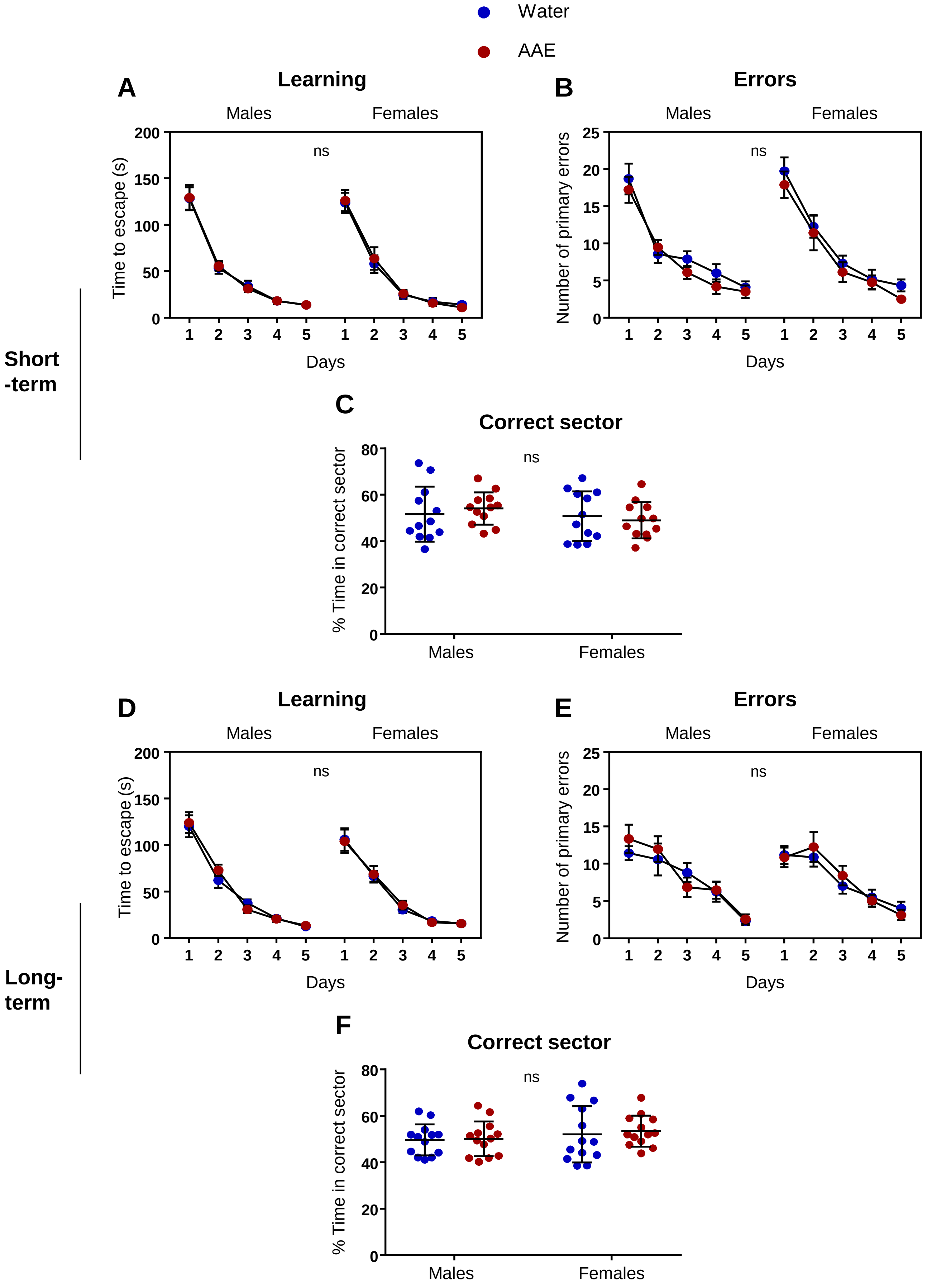

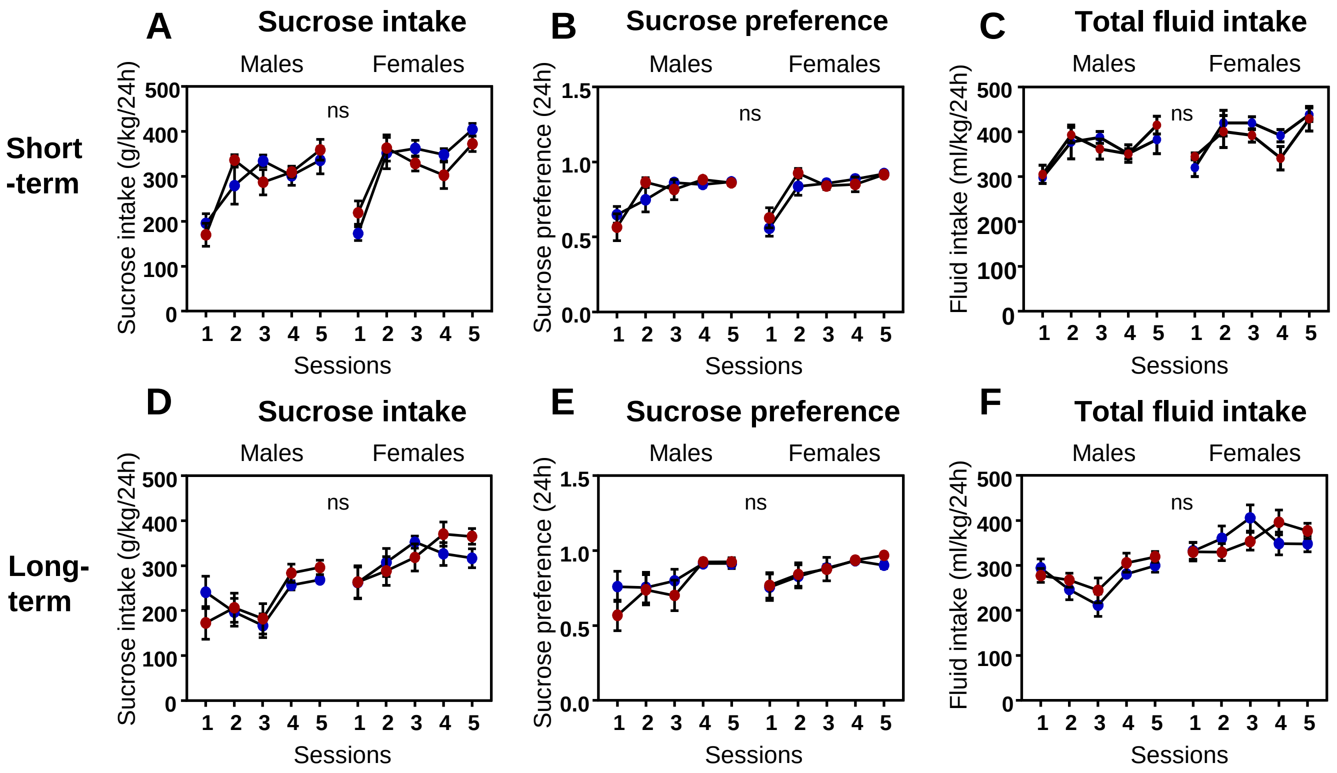

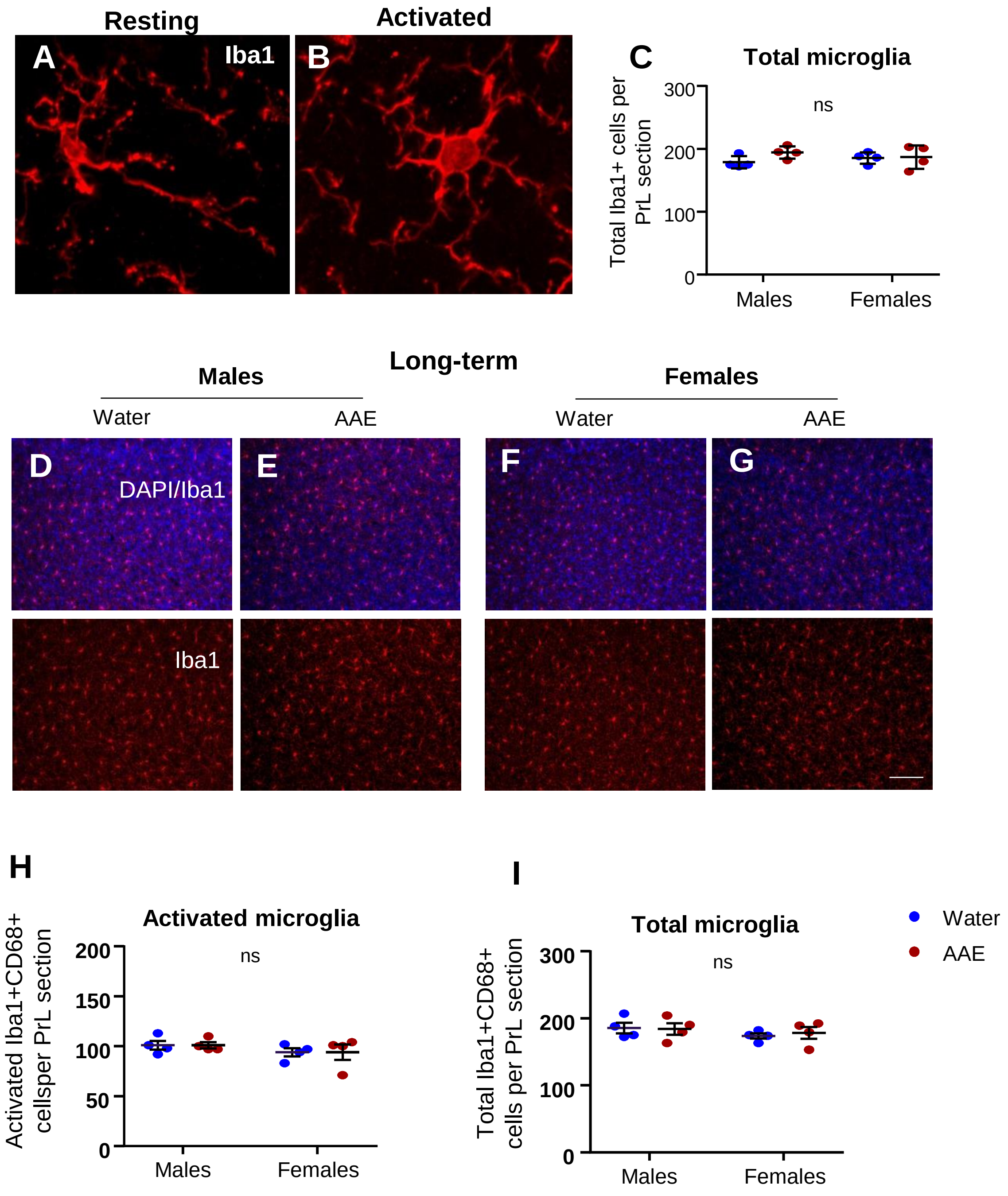
